# ATR2^Cala2^ from *Arabidopsis*-infecting downy mildew requires 4 TIR-NLR immune receptors for full recognition

**DOI:** 10.1101/2023.04.25.538220

**Authors:** Dae Sung Kim, Alison Woods-Tör, Volkan Cevik, Oliver J. Furzer, Yufei Li, Wenbo Ma, Mahmut Tör, Jonathan D. G. Jones

## Abstract

- *Arabidopsis* Col-0 RPP2A and RPP2B confer recognition of *Arabidopsis* downy mildew (*Hyaloperonospora arabidopsidis* [*Hpa*]) isolate Cala2, but the identity of the recognized ATR2^Cala2^ effector was unknown.
- To reveal *ATR2^Cala2^*, an F_2_ population was generated from a cross between *Hpa*-Cala2 and *Hpa*-Noks1. We identified ATR2^Cala2^ as a non-canonical RxLR-type effector that carries a signal peptide, a dEER motif, and WY domains but no RxLR motif. Recognition of *ATR2^Cala2^*and its effector function were verified by biolistic bombardment, ectopic expression and *Hpa* infection.
- ATR2^Cala2^ is recognized in accession Col-0 but not in Ler-0 in which RPP2A and RPP2B are absent. In *ATR2^Emoy2^* and *ATR2^Noks1^* alleles, a frameshift results in an early stop codon. RPP2A and RPP2B are essential for the recognition of ATR2^Cala2^. Stable and transient expression of *ATR2^Cala2^* under 35S promoter in *Arabidopsis* and *Nicotiana benthamiana* enhances disease susceptibility.
- Two additional Col-0 TIR-NLR (TNL) genes (*RPP2C* and *RPP2D*) adjacent to *RPP2A* and *RPP2B* are quantitatively required for full resistance to *Hpa*-Cala2.
- We compared *RPP2* haplotypes in multiple *Arabidopsis* accessions and showed that all 4 genes are present in all ATR2^Cala2^-recognizing accessions.

## Introduction

Plants, like animals, are constantly exposed to potentially damaging pathogens, and like invertebrates but unlike mammals, rely solely on innate immunity (Jones and Takemoto, 2004). The plant immune response is highly effective but must be activated early to thwart pathogens, and activation requires detection of pathogen molecules by cell surface and intracellular immune receptors. Cell-surface receptors usually detect relatively conserved pathogen-associated molecular patterns (PAMPs) and activate pattern-triggered immunity (PTI) (Monaghan and Zipfel, 2012; Boutrot and Zipfel, 2017). During plant-microbe co-evolution, pathogens evolved the ability to deliver effector proteins to host cells that suppress PTI, enabling pathogen growth (Feng and Zhou, 2012). In turn, plants evolved intracellular immune receptors, often encoded by resistance (*R*) genes, that either directly or indirectly detect the presence of pathogen effector proteins (Nürnberger et al., 2004; Chisholm et al, 2006; Jones and Dangl, 2006; Jones et al., 2016) and activate effector-triggered immunity (ETI) (Dodds and Rathjen, 2010; Dangl et al., 2013). Recognized effectors are for historical reasons often referred to as avirulence (Avr) proteins. Intracellular recognition usually requires nucleotide-binding, leucine-rich repeat (NB-LRR, or NLR) immune receptors. NLR activation results in an elevated immune response, characterized by generation of reactive oxygen species, cell wall fortification, activation of defense-associated genes, and a localized cell death known as the hypersensitive response (HR) (Spoel and Dong, 2012). Many cases of matching *R* and *Avr* genes have been described (Jones and Dangl, 2006; Bernoux et al., 2011). However, in some examples, disease resistance against a pathogen isolate or recognition of an Avr protein, requires the coordinate function of pairs of NLR genes (Eitas and Dangl, 2010). Recent detailed studies on the *Arabidopsis* TIR-NLR pair RRS1 and RPS4, and the rice CC-NLR pairs RGA4/RGA5 and Pik-1/Pik-2 reveal how such protein pairs function together. The paired partners interact physically to form a receptor complex in which each protein plays distinct roles in effector recognition or signalling activation, exemplifying a conserved mode of action of NLR pairs in diverse plants (Cesari et al., 2014; Sarris et al., 2015; Ma et al., 2018). Such gene pairs are often divergently transcribed. Interestingly, 10 of 11 pairs of TIR-NLR genes show a head-to-head configuration in *Arabidopsis* (Meyers et al., 2003). Divergent transcription may assure balanced levels of the protein pair to meet a strict stoichiometric requirement to act together, possibly in a complex (Narusaka et al., 2009). However, the *Arabidopsis RPP2* locus that confers resistance to downy mildew (Sinapidou et al., 2004), comprises two genes, *RPP2A* and *RPP2B* that are not divergently transcribed.

Downy mildews are obligate biotrophic oomycete pathogens and can cause severe diseases on many different vegetable crops (Thines and Kamoun, 2010; Tör et al., 2023). *Hyaloperonospora brassicae* causes severe disease in Chinese cabbage (*Brassica rapa* L. ssp. *pekinensis*), which is native to China and is one of the most important vegetables in Asia. In epidemic seasons with warm temperatures and high humidity, 80%-90% of Chinese cabbage plants are infected by *H. brassicae*, leading to a 30%-50% reduction of production (Li et al., 2011). Downy mildew caused by *Bremia lactucae* is the most important disease in lettuce (*Lactuca sativa* L.) reducing yield and decreasing the quality of the marketable portion (Parra et al., 2021). Downy mildew caused by *Plasmopara viticola* can lead to severe damage to grapevines (Li et al., 2015). Cucumber (*Cucumis sativus* L.) downy mildew, caused by *Pseudoperonospora cubensis*, is a major destructive and widespread disease of cucumber plants (Zhang et al., 2019). There has been an increasing interest in the molecular mechanisms of downy mildew resistance (Liu et al., 2021). The use of cultivars carrying dominant resistant (*Dm*) genes in lettuce is the most effective way to control downy mildew caused by *B. lactucae* (Parra et al., 2021). The model plant *Arabidopsis* is susceptible to the downy mildew *Hyaloperonospora arabidopsidis (Hpa)* (Slusarenko and Schlaich, 2003). Various *RPP* (*<u>R</u>esistance to <u>P</u>eronospora parasitica,* the former name of *Hpa)* genes in different accessions confer resistance to specific *Hpa* isolates (Asai et al., 2018).

Obligate biotrophic pathogen races or isolates differ in their capacity to evade or suppress host recognition (Oliver and Ipcho, 2004). Oomycete pathogens deploy effector proteins, with a signal peptide and typically the signature amino acid motifs RxLR and DEER (Rehmany et al., 2005). A subset of such effectors also carries a variable number of repeats of a WY domain (Win et al., 2012). RxLR effectors has been intensively investigated since their discovery (Anderson et al., 2015; Wood et al., 2020).

The *Arabidopsis*/*Hpa* pathosystem reveals extensive genetic diversity in host *Resistance* (*RPP*) and cognate pathogen *ATR* (*<u>A</u>rabidopsis thaliana* recognized) genes (Coates and Beynon, 2010; Asai et al., 2018). Using an *Hpa* reference genome (Baxter et al., 2010), 475 *Hpa* gene models were identified that encode effector candidates in *Hpa*-Emoy2, using the following criteria: (1) proteins with a signal peptide and canonical RxLR motif, like ATR1, ATR13, and ATR39 (HaRxLs) (Rehmany et al., 2005; Allen et al., 2004; Goritschnig et al., 2012), (2) RxLR-like proteins with at least one non-canonical feature, like ATR5 (HaRxLLs) (Bailey et al., 2011), (3) putative Crinkler-like proteins with RxLR motif (HaRxLCRNs) (Win et al., 2007), (4) homologous proteins based on amino acid sequence similarity over the N-terminal region including a signal peptide and RxLR motif (e.g., HaRxL1b, HaRxLL2b, and HaRxLCRN3b) (Asai et al., 2014).

Several *RPP* genes, including *RPP1*, *RPP2A* and *RPP2B*, *RPP4*, *RPP5*, *RPP8*, *RPP13*, and *RPP39* encode NLR immune receptors (Holub 2008). Genetic analyses of avirulence in *Hpa* has confirmed a gene-for-gene relationship for *ATR* genes (Holub et al. 1994) with their corresponding *RPP* genes. Recognized *Hpa* effectors ATR1, AvrRPP4, ATR5, ATR13 and ATR39 have been identified for RPP1, RPP4, RPP5, RPP13, and RPP39 (Allen et al., 2004; Rehmany et al., 2005; Bailey et al., 2011; Goritschnig et al., 2012; Asai et al., 2018). For example, the *RPP1* locus, which contains a complex resistance gene cluster, was originally identified in *Arabidopsis* accession Wassilewskija (Ws-2) (Botella et al., 1998). Several members of the *RPP1* gene family confer resistance against isolates of *Hpa* (Botella et al., 1998; Rehmany et al., 2005; Sohn et al., 2007), including *RPP1*-WsA, *RPP1*-WsB, *RPP1*-WsC, and *RPP1*-NdA, while *RPP1*-like genes from other accessions have been implicated in hybrid incompatibility (Bomblies et al., 2007). Proteins encoded by two *RPP1* alleles have been shown to recognize the cognate effector ATR1 from *Hpa* (Rehmany et al., 2005; Krasileva et al., 2010; Ma et al., 2020). The R proteins RPP1-WsB and RPP1-NdA share a common TNL domain architecture and are 87% identical at the amino acid level. Although polymorphisms are present throughout their coding sequences, most of the differences occur in the LRR region and include both single amino acid polymorphisms and short insertions and deletions. ATR1 belongs to a simple locus in *Hpa* with allelic variants present in different pathogen races (Rehmany et al., 2005; Krasileva et al., 2010). ATR1 carries an N-terminal eukaryotic signal peptide and an RxLR motif (Rehmany et al., 2005; Birch et al., 2006) and associates with its cognate RPP1 immune receptor via its LRR domain (Krasileva et al., 2010). The tetrameric complex containing four RPP1 and four ATR1 molecules is mediated by direct binding of ATR1 to a C-terminal jelly roll/Ig-like domain (C-JID) and the LRRs of RPP1 (Ma et al., 2020).

RPP2A and RPP2B are both required for resistance to *Hpa* isolate Cala2 (Sinapidou et al., 2004), but their cognate effector ATR2 was not identified previously. Adjacent to *RPP2A* (*At4g19500*) and *RPP2B* (*At4g19510*) (Sinapidou et al., 2004) lie two other TNL encoding genes (*At4g19520* and *At4g19530*, hereafter *RPP2C* and *RPP2D*). They comprise, in a head-to-head conformation, a similar gene pair to RRS1 and RPS4, including a C-terminal extended post-LRR domain. In this research, we aimed to clone and characterize *ATR2*, investigate its virulence function and its contribution to effector recognition by the four genes at the *RPP2* locus.

Using an F_2_ population generated from a cross between *Hpa*-Cala2 and *Hpa*-Noks1 (Bailey et al, 2011), we positionally cloned *ATR2*. We show here that functional ATR2 is absent from the reference Emoy2 genome and its annotated proteome, that ATR2 confers elevated disease susceptibility when expressed *in planta* and that all four RPP2 paralogs contribute to its full recognition.

## Materials and Methods

### Plant materials and growth

*Arabidopsis* accessions, Col-0, Ler-0, Oy-0, Ws-2, Ws-2 *eds1* and CW84, which is an *Hpa* susceptible recombinant inbred line generated from a cross between Col-0 and Ws-2 (Botella et al., 1998) were grown at 22°C under short-day condition (10 h light/14 h dark) and *Nicotiana benthamiana* plants were grown at 25 °C under a 16-h photoperiod and an 8-h dark period in environmentally controlled growth cabinets.

### Positional cloning of *ATR2^Cala2^*

The crossing of *Hpa*-Cala2 and *Hpa*-Noks1 and production of F_2_ mapping population from a single-spored CaNo F1 were described previously (Bailey et al., 2011). Initially, segregating 52 random CaNo F_2_ isolates were bulked up on Ws-*eds1* seedlings and tested on Col-5 to determine the genetic nature of *ATR2*. As the genomic sequences of parental isolates were not available then, a similar approach to clone *ATR5* (Bailey et al., 2011) was taken where DNA was isolated from individual CaNo F_2_ isolates and a bulk segregant analysis was employed to clone *ATR2*. Two different bulks were constructed from the CaNo F_2_ individuals (18 F_2_s with *ATR2*/± and 17 with *atr2/atr2* genotypes) and AFLP was carried out with *EcoR*I and *Mse*I primer pairs as described (Bailey et al., 2011). Fifteen polymorphic AFLP fragments were identified and converted to CAPS markers to map *ATR2* onto publicly available BAC contigs. As the *Hpa*-Emoy2 reference genome became available, we used these markers to identify the *Hpa*-Emoy2 SuperContig9. As the number of recombinants were very low, additional CaNo F_2_ isolates were generated and Illumina paired-end sequencing data of CaNo F_2_ bulks were obtained. As the genomic data for *Hpa*-Cala2 and *Hpa*-Noks1 became available (Woods-Tör et al., 2018), the bulk sequences were mapped onto *Hpa*-Cala2 genome as described (Woods-Tör et al., 2018) and SNP markers were identified within the interval. Further markers were generated from the identified SNP sites and using a total of 130 CaNo F_2_ isolates, we mapped *ATR2* to a 186.5 kb interval on *Hpa*-Cala2 SuperContig9. Further markers were generated and new F_2_ isolates were obtained, and the locus was mapped to a 112 kb interval. We compared genomic sequences of *Hpa*-Emoy2, *Hpa*-Noks1and *Hpa*-Cala2 for the interval to identify possible candidates for *ATR2*. All PCR amplifications for mapping were performed as described (Woods-Tör et al., 2018).

### Pathogen assays

*Hpa* isolates, *Hpa*-Emoy2, *Hpa*-Noks1 and *Hpa*-Cala2 were propagated and maintained by weekly sub-culture on 14-day-old *Arabidopsis* seedlings. Preparation of inoculum for experiments, and the assessment of sporulation were as described previously in Bailey et al., 2011.

*Pseudomonas syringae* pv. *tomato* (*Pst*) DC3000 was grown in King’s B broth (10 g peptone, 15 g glycerol, 1.5 g K_2_HPO_4_ and 5 mM MgSO_4_ per litre) containing 50 μg ml^-1^ rifampicin. Leaves of 5-week-old *Arabidopsis* plants were infiltrated with 10^5^ cfu ml^-1^ of *Pst* DC3000 using a needleless syringe. Bacterial growth was measured at 0- and 3-days post inoculation (dpi).

*Phytophthora infestans* isolate 88069 was grown on Rye Agar at 19°C for 2 weeks. Plates were flooded with 5 ml cold H_2_O and scraped with a glass rod to release zoospores. The resulting solution was collected in a falcon tube and zoospore numbers were counted using a hemacytometer and adjusted to 2 X 10^4^ zoospores/ml and 10 µl droplets were inoculated onto the abaxial side of leaves of intact *N. benthamiana* plants. Inoculated leaves were then stored on moist tissues in sealed boxes.

### Plasmid construction

All the constructs used in this study were generated using USER (Uracil-Specific Excision Reagent) enzyme cloning method (Geu-Flores et al. 2007). Briefly, target DNA to be cloned into destination USER vectors, pICSLUS0003 or pICSLUS0004 (archived in TSL Synbio) was amplified using *PfuTurbo® C_x_* polymerase (Agilent Technologies) with uracil-containing primer pair then assembled with desired tag (“Hellfire” including 6-His and 3-FLAG epitopes), linearized vector and USER enzyme (NEB). For transient gene expression in *N. benthamiana* or *N. tabacum*, *ATR2* candidates without signal peptide were cloned and assembled.

### Bombardment and luciferase assays

Co-bombardment assays were performed as described previously with some modifications (Bailey et al., 2011). Briefly, *Arabidopsis* plants were grown with short-day condition until 6 weeks old. Detached leaves were placed on a 1% MS agar in a petri dish. One µm of tungsten particles were coated with the plasmids carrying genes *ATR2* and luciferase under 35S promoter. Bombardments were performed using a Bio-Rad PDS-1000 (He) apparatus with 1,100 p.s.i. rupture disks, as per manufacturer’s instructions. For each replicate, a leaf from both test and control plant genotypes were co-bombarded together in a single shot. Bombarded leaves were put into 10 ml plastic vials filled with water 1 cm from the bottom and were incubated at 25°C for 20 h.

For the luciferase assay a Dual Reporter Luciferase Assay system (Promega) was used. Four transiently bombarded leaf events were pooled together and crushed in Luciferase Cell Culture Lysis buffer (Promega). The extract was centrifuged at 12,000 rpm for 10 min at 4 °C. 20 µl of the lysate was then dispersed in 96 well plate in triplicates and analyzed on Varioskan Flash Instrument by injecting 100 µl of luciferase assay reagent II, which includes substrate and reaction buffer. A 10 second read time was used to measure luciferase activity for each well.

### Expression analysis

Total RNA was isolated from three biological replicates using the RNeasy Plant Mini Kit (Qiagen) with the Dnase treatment (Qiagen). cDNA was synthesised using SuperScript IV Reverse Transcriptase (ThermoFisher). For *ATR2* gene expression analysis during *Hpa* infection, reverse transcription (RT)-PCR was performed.

### Transient expression in *Nicotiana* species

*Agrobacterium tumefaciens* GV3101 strain harbouring *ATR2* candidate fused to 35S promoter was streaked on selective media and incubated at 28 °C for 24 hours. A single colony from the streaked inoculum was transferred to liquid LB media with appropriate antibiotic and incubated at 28 °C for 48 hours in a shaking incubator at 180 rpm. The cultures were centrifuged at 3,000 rpm for 5 min and resuspended in infiltration buffer (10 mM MgCl2, 10 mM MES, pH 5.7) and acetosyringone was added to a final concentration of 200 μM at OD_600_ of 1.0. The abaxial surface of 4-weeks old *N. tabacum* or *N. benthamiana* was infiltrated with a 1 ml needleless syringe (Kim et al., 2015).

### *Arabidopsis* transformation

*Arabidopsis* accessions Col-0 and Ler-0 expressing *ATR2* candidate gene, and CW84 expressing Col-*RPP2* cluster harbouring JAtY clone (Zhou et al., 2011) were transformed using *A. tumefaciens* strain GV3101 by flower dipping method (Clough and Bent, 1998).

### RPP2 cluster haplotype analyses

Full-length amino acid sequences of individual RPP2A, RPP2B, RPP2C and RPP2D from 64 different *Arabidopsis* accessions were extracted from pan-NLRome data (Van de Weyer et al., 2019). Each group of RPP2 was aligned to each other using Geneious Prime software to investigate haplotype patterns of RPP2 clusters. Pfam (Punta et al., 2012; http://pfam-legacy.xfam.org) was used for domain analysis in RPP2 cluster.

### Protein structure modelling

Protein tertiary structure model of full-length ATR2^Cala2^ was generated by Alphafold 2 (Jumper et al., 2021; Varadi et al., 2021). The region spanning the Y-WY sequences was extracted and superimposed with the structure of full-length PsPSR2 using PyMOL Molecular Graphics System, Version 1.2r3pre, LLC (Xiong et al., 2014; He et al., 2019; Hou et al., 2019). Secondary structures and surface accessibility of ATR2^Cala2^ were predicted by NetSurfP-3.0 (Høie et al., 2022). Alignment with published LWY effectors revealed the conserved W and Y residues in ATR2^Cala2^ and the corresponding Y and WY modules (He et al., 2019).

### Accessions

Genomic sequences of parental isolates can be found under accession numbers GCA_001414265.1 for *Hpa*-Cala2, GCA_001414525.1 for *Hpa*-Noks1, GCA_000173235.2 for *Hpa*-Emoy2 in NCBI. The raw sequence reads from the genomics sequencing of bulks are available from the Sequenced Read Archive (SRA) under accession numbers SRX13788375 (avirulent) and SRX13788374 (virulent). *ATR2^Cala2^* and *ATR2^Emoy2^* sequences were deposited in NCBI (GenBank accession no. ON994189 and ON994190, respectively). Resistance (*R*) gene sequence capture (RenSeq) raw sequencing data of FN2 (*rpp2a-1*) mutant is available from the SRA (accession no. PRJNA955397).

## Results

### Mapping *ATR2*

Positional cloning was used to identify the *ATR2* locus in *Hpa*-Cala2. A segregating CaNo F_2_ population (Bailey et al., 2011) was used to define the *ATR2* locus. Initially, 52 randomly chosen F_2_ isolates were tested on Col-5. A single semi-dominant avirulence determinant designated *ATR2^Cala2^* segregated in the F_2_ population (avirulence:virulence ratio was 40:12, with chi-square = 0.1025 and P = 0.74, Table S1). Bulked segregant analysis was used to identify AFLP markers that are linked to *ATR2* in the CaNo F_2_ population. AFLP markers were cloned and converted to CAPS markers, which were then used for mapping *ATR2*. The reference genome of *Hpa-*Emoy2 was still being generated during this early mapping work and genomic sequence data for *Hpa*-Cala2 and *Hpa*-Noks1 were not available. We screened an *Hpa*-Emoy2 BAC library (Rehmany et al., 2003) with the *ATR2-*linked CAPS markers. Several BACs were identified, and BAC-end sequences were used to assemble a small contig around the *ATR2* locus with markers that revealed one recombinant from one side and that co-segregate on the other side with a gap in the middle (Fig. S1), making it difficult to narrow the *ATR2* locus onto a single BAC (Fig. S1).

Once *Hpa*-Emoy2 genomic sequence became available, we transferred the AFLP-derived CAPS markers to *Hpa*-Emoy2 SuperContig9, which helped us to identify the physical location of the locus (Fig. S1, S2).

We then generated 100 bp paired-end Illumina HiSeq2500 sequencing data from the two newly bulked (virulent and avirulent) pools, comprising 110 million reads for the virulent bulk and 104 million reads for the avirulent bulk. We also utilized *Hpa*-Cala2 and *Hpa*-Noks1 genomic sequences (Woods-Tör et al., 2018) to identify SNPs between *Hpa*-Cala2 and *Hpa*-Noks1 genomes. Using new markers generated from these SNPs, we established an interval of 186.5kb on *Hpa*-Emoy2 SuperContig9: 656515-843042 (Fig. S1). We generated further markers and F_2_ isolates and narrowed the *ATR2* locus to a 112 kb interval on *Hpa*-Emoy2 SuperContig9: 708503-820527 (Table 1 and Table S2).

**Table 1.**
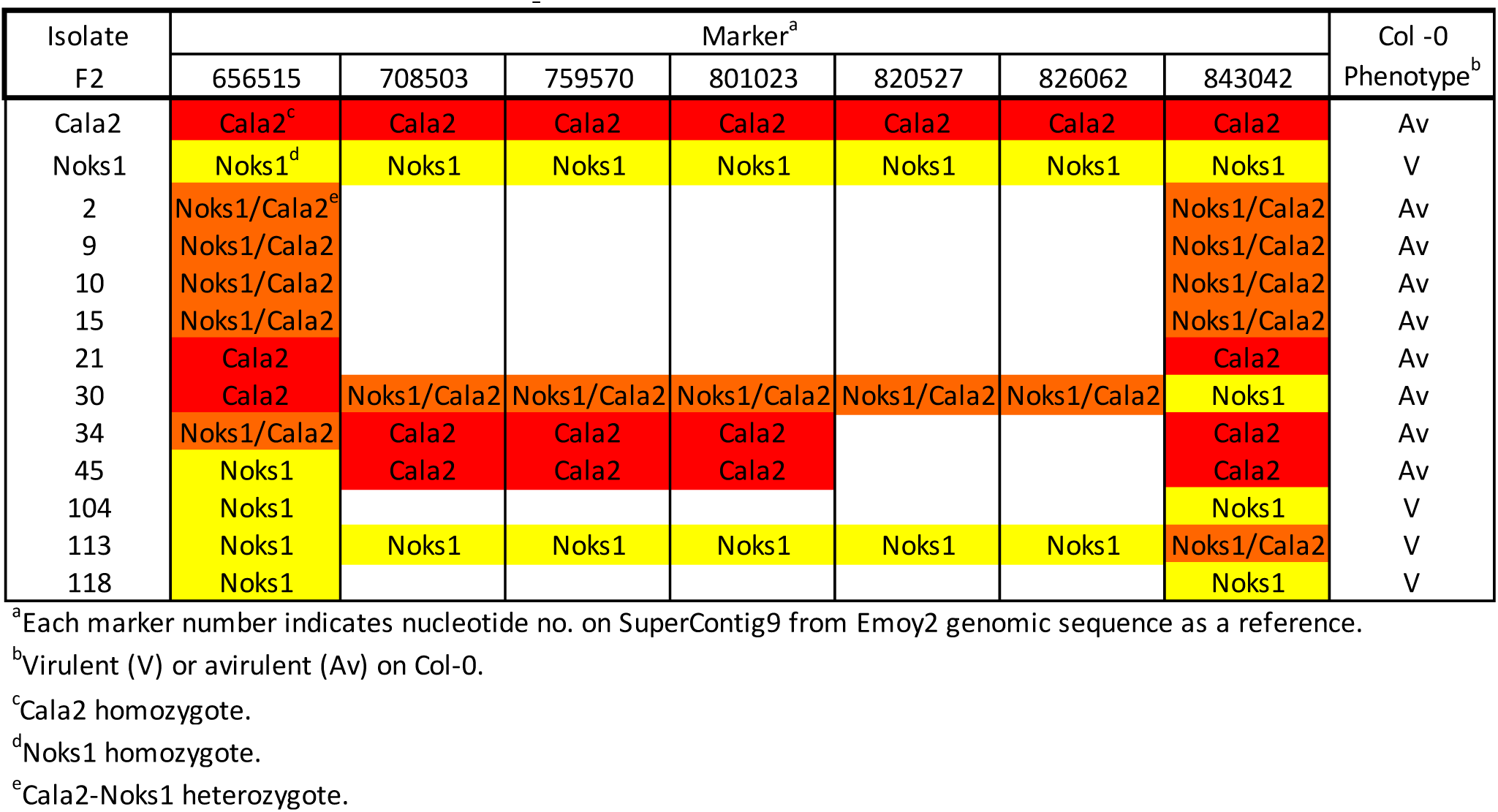
Intervals of *ATR2* from F2 CaNo F_2_ isolates.

| Isolate<br>F2 | Marker<sup>a</sup> |  |  |  |  |  |  | Col -0<br>Phenotype<sup>b</sup> |
| --- | --- | --- | --- | --- | --- | --- | --- | --- |
|  | 656515 | 708503 | 759570 | 801023 | 820527 | 826062 | 843042 |  |
| Cala2 | Cala2<sup>c</sup> | Cala2 | Cala2 | Cala2 | Cala2 | Cala2 | Cala2 | Av |
| Noks1 | Noks1<sup>d</sup> | Noks1 | Noks1 | Noks1 | Noks1 | Noks1 | Noks1 | V |
| 2 | Noks1/Cala2<sup>e</sup> |  |  |  |  |  | Noks1/Cala2 | Av |
| 9 | Noks1/Cala2 |  |  |  |  |  | Noks1/Cala2 | Av |
| 10 | Noks1/Cala2 |  |  |  |  |  | Noks1/Cala2 | Av |
| 15 | Noks1/Cala2 |  |  |  |  |  | Noks1/Cala2 | Av |
| 21 | Cala2 |  |  |  |  |  | Cala2 | Av |
| 30 | Cala2 | Noks1/Cala2 | Noks1/Cala2 | Noks1/Cala2 | Noks1/Cala2 | Noks1/Cala2 | Noks1 | Av |
| 34 | Noks1/Cala2 | Cala2 | Cala2 | Cala2 |  |  | Cala2 | Av |
| 45 | Noks1 | Cala2 | Cala2 | Cala2 |  |  | Cala2 | Av |
| 104 | Noks1 |  |  |  |  |  | Noks1 | V |
| 113 | Noks1 | Noks1 | Noks1 | Noks1 | Noks1 | Noks1 | Noks1/Cala2 | V |
| 118 | Noks1 |  |  |  |  |  | Noks1 | V |

<sup>a</sup>Each marker number indicates nucleotide no. on SuperContig9 from Emoy2 genomic sequence as a reference.
<sup>b</sup>Virulent (V) or avirulent (Av) on Col-0.
<sup>c</sup>Cala2 homozygote.
<sup>d</sup>Noks1 homozygote.
<sup>e</sup>Cala2-Noks1 heterozygote.

### Genes in the *ATR2* interval encode effector-like proteins

We then used comparative genomics for this region using *Hpa*-Cala2, *Hpa*-Noks1 and *Hpa*-Emoy2 contigs, and found three putative candidate effector genes (*A2C1*, *A2C2* and *A2C3,* for *ATR2* candidates 1, 2 or 3) (Fig. S2). *A2C1* and *A2C2* correspond to RxLL457, and another RxLR protein, respectively (Fig. S3a, b, c). *A2C3* was predicted to encode a non-canonical RXLR (GHVR) protein with dEER and WY motifs (Fig. 2a).

Recognition of these candidates by RPP2 was evaluated by biolistic co-bombardment of 35S:*A2C1, 2 or 3* constructs with 35S:*Luciferase* and assessing luciferase eclipse. If localized cell-death is initiated upon ATR2^Cala2^ recognition, luciferase activity is compromised compared to leaf tissues expressing luciferase only. An *Hpa-*Cala2-susceptible recombinant inbred line *Arabidopsis* CW84 (Bailey et al., 2011) was used as a control. CCG28, an oomycete *Albugo candida* effector which is recognized by Col-0 and CW84 was also used as a positive control (Redkar et al., 2023). Reduced luciferase activity was observed in *CCG28*-co-bombarded tissues, but no reduction was detected when either *A2C1^Noks1^* or *A2C1^Cala2^* alleles or *A2C2^Emoy2^*or *A2C2^Cala2^* alleles were co-bombarded, indicating they are not recognized in Col-0 or in CW84. Thus, neither of them is *ATR2^Cala2^* (Fig. S3d). When *A2C1* was transiently expressed with *RPP2A* and/or *RPP2B* on *N. benthamiana* or tobacco leaves, no HR was observed while *AvrRPP4*-*RPP4* combination triggered strong HR, acting as a positive control and confirming our bombardment assays (Fig. S4a). Protein expression of epitope-tagged A2C1 and A2C2 was verified by protein gel blot (Fig. S4a). Similarly, neither of *A2C2* alleles from *Hpa*-Emoy2, Cala2 or Noks1 triggered cell death in *N. benthamiana* by transient expression alone or with *RPP2A* and/or *RPP2B* (Fig. S4b).

### *A2C3^Cala2^* encodes an RxLR effector-like candidate for *ATR2^Cala2^*

*A2C3* was identified in *Hpa*-Cala2 after re-sequencing of the 5 kb upstream of *A2C1* and *A2C2,* which includes a highly polymorphic region of Cala2 compared to Emoy2 (Fig. 1a). We found a transposable element in this region of the *Hpa*-Emoy2 genome and a 2.3 kb deletion in *Hpa*-Cala2. We also found a cytosine insertion on the *Hpa*-*802071* coding region in Cala2 which created a frameshift in the *Hpa-802071* coding region (Fig. 1b). To determine whether *A2C3^Cala2^* co-segregates with recognition by *RPP2* in the F_2_ population, *A2C3* alleles were amplified and sequenced from 12 CaNo F_2_ segregants, *Hpa*-Emoy2, *Hpa*-Noks1 and *Hpa*-Cala2 (Fig. S5). All avirulent F_2_s were homozygous- or heterozygous for *A2C3^Cala2^,* while all virulent F_2_s were homozygous for *A2C3^Emoy2/Noks1^* (Fig. 1c, S5c). Of *Hpa*-Emoy2, *Hpa*-Noks1 and *Hpa*-Cala2, only *Hpa*-Cala2 carries this non-canonical RxLR effector candidate (A2C3^Cala2^), which has a signal peptide, a dEER, and Y and WY motifs (Fig. 2a, S6a). *A2C3^Emoy2^* and *A2C3^Noks1^*alleles were identical to each other with early stop codons caused by a frameshift. These data suggested *A2C3^Cala2^* might be *ATR2^Cala2^*. We analysed synonymous and non-synonymous SNPs in *A2C3* alleles among 7 different *Hpa* isolates for which genomic data are available. Many non-synonymous SNPs are found only in *Hpa*-Cala2, indicating specificity of *A2C3^Cala2^* (Table S3). Alignment of *A2C3^Cala2^* with *Phytophthora* LWY effectors revealed conserved W and Y residues and the corresponding Y and WY modules of A2C3^Cala2^ (He et al., 2019). The Y-WY modules of A2C3^Cala2^ in an A2C3^Cala2^ structural model predicted by Alphafold 2 were extracted and superimposed with PsPSR2 which is a typical RxLR effector with one WY motif and six LWY motifs. This region was well matched on PsPSR2 Y5-LWY6 (region from Y5 of LWY5 to LWY6) with Root Mean Square Deviation (RMSD) = 2.305 (Fig. 2b). This structural comparison also revealed that there is an L-like module between Y and WY modules of A2C3^Cala2^ even though this was not predicted by amino acid sequence comparison (Fig. 2b, S6b).

**Figure 1.**
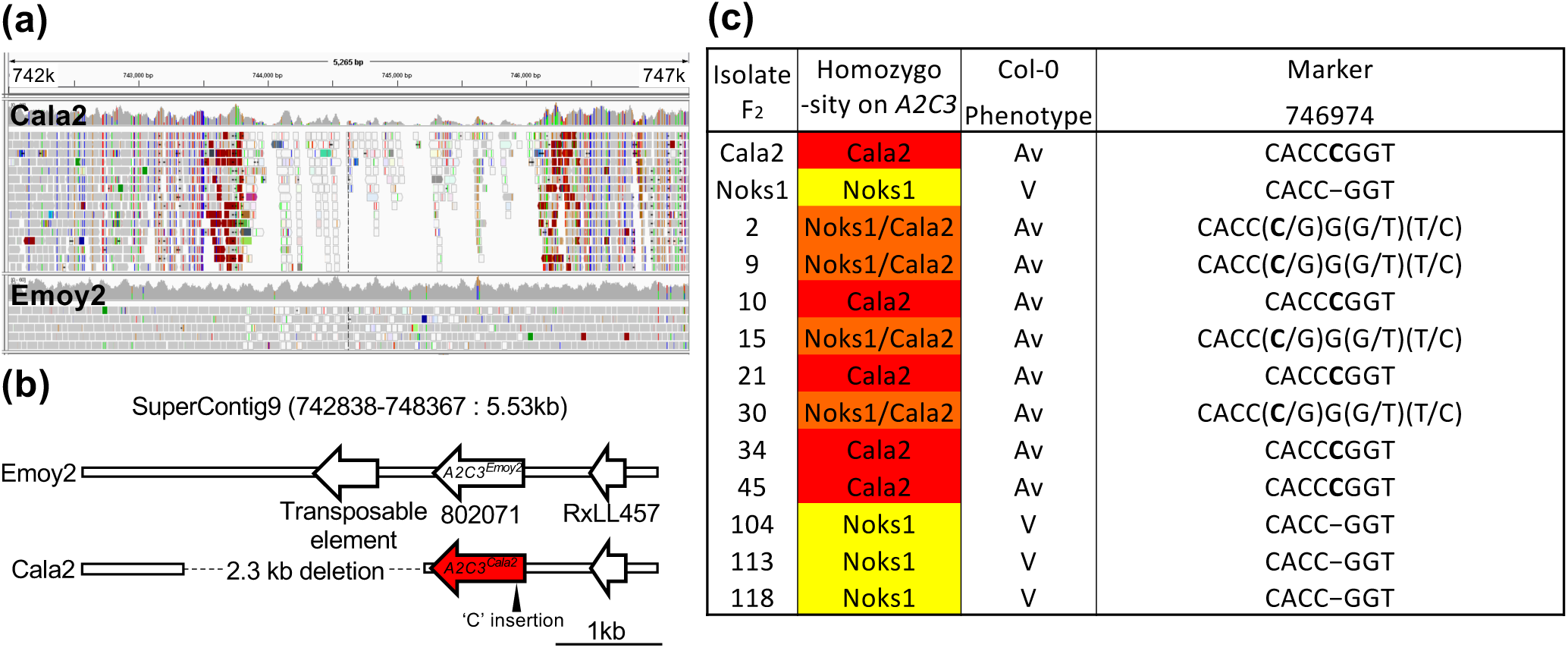
Genetic determination of *A2C3^Cala2^* co-segregation from avirulent F_2_ isolates. (a) Comparison of highly-polymorphic region of Cala2 with Emoy2 from 742k to 747k of SuperContig9. Screenshot was captured from IGV software. (b) Transposable element next to ‘802071’ on Emoy2 and 2.3kb deletion on Cala2 of assigned region. *A2C3^cala2^* allele is highlighted with red colour. A cytosine (C) insertion at 746974 indicated with a black triangle. (c) Analyses of homo- or heterozygosity, avirulent (Av) or virulent (V) on Col-0 and a segregated cytosine (C) insertion in Av isolates at the frameshift region (746974) by sequencing from Fig. S5. Red: Cala2 homozygote; Yellow: Noks1 homozygote; Orange: Cala2-Noks1 heterozygote on *A2C3*. Inserted cytosine was bold highlighted.

**Figure 2.**
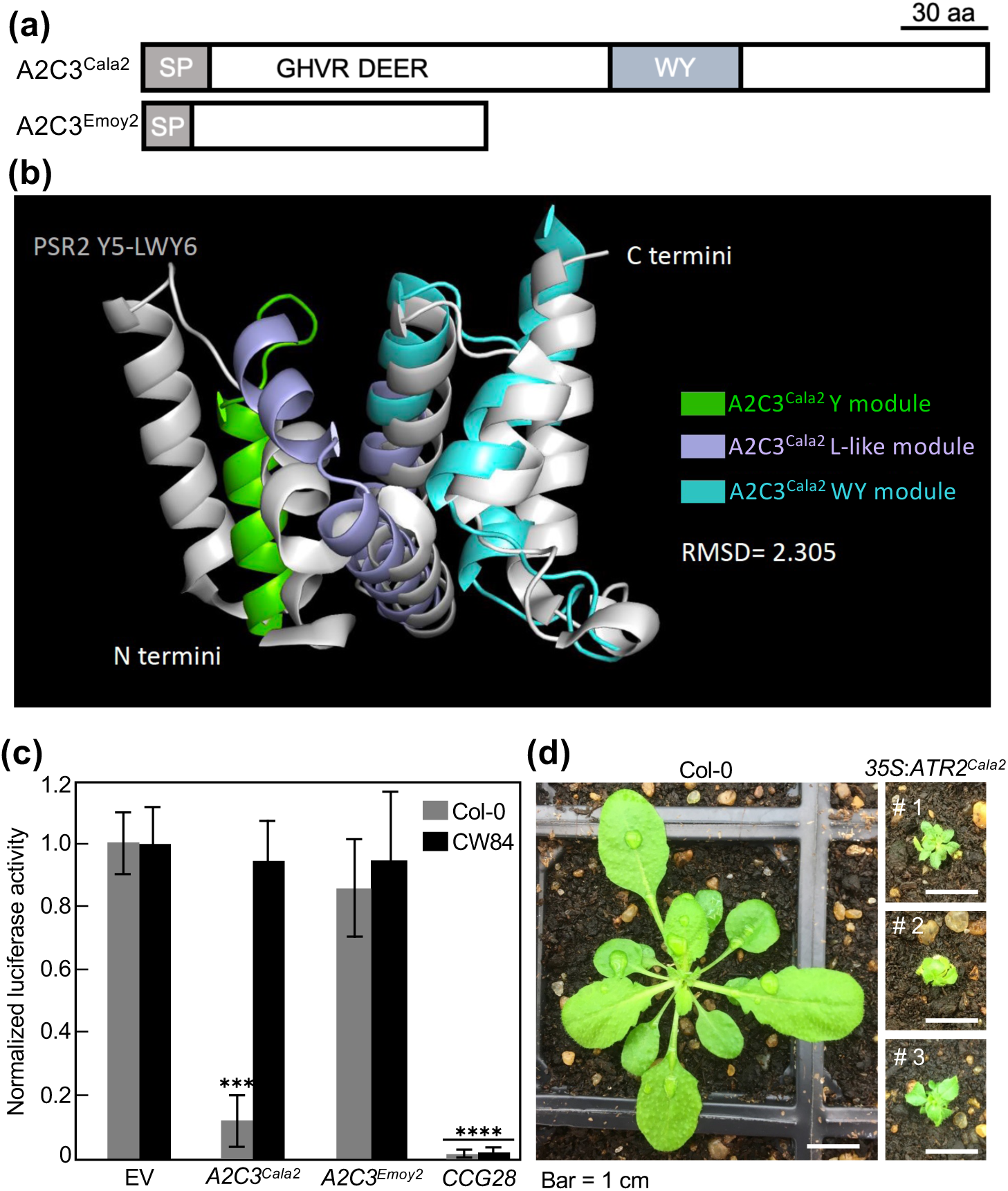
A2C3^Cala2^ recognition capacity in Col-0. (a) Schematic diagrams of A2C3 of *Hpa* Cala2 and Emoy2. (b) Alphafold 2 prediction of Y and WY modules of A2C3^Cala2^, and super-imposition with PsPSR2. PsPSR2 contains seven (L)WY units with Y5-LWY6 showing the highest similarity with ATR2^Cala2^. This structural comparison also revealed an “L” like fold between the “Y” and “WY” sequence in ATR2^Cala2^. (c) Biolistic bombardment of *A2C3* with luciferase into Col-0 and CW84 in which RPP2 is absent. Data are mean ± standard deviations from three independent experiments). Asterisks (***, P < 0.001; ****, P < 0.0001) indicate statistical significance compared with luciferase alone in Col-0 or CW84 by two-way ANOVA with Tukey’s multiple comparison test. EV, empty vector. (d) Transgenic Arabidopsis Col-0 expressing *ATR2^Cala2^* under 35S promoter. Bar = 1 cm.

We determined the expression of *A2C3^Cala2^* alleles during *Hpa* infection. *Arabidopsis* Oy-0 accession was infected with *Hpa*-Emoy2 and Ler-0 was infected with *Hpa*-Cala2, and Col-*eds1* was used as hyper-susceptible control. *A2C3^Cala2^* is expressed at 3 dpi (Fig. S7a). Previously, Asai et al. (2014) performed expression profiling of *Hpa* genes from *Hpa* Emoy2. RNA-seq data of *Hpa*-*802071* (*A2C3^Emoy2/Noks1^*) were retrieved from the data. Again, *A2C3^Emoy2/Noks1^* is induced after infection and shows the highest expression at 3 dpi (Fig. S7b). We proceeded to further evaluate the *A2C3^Cala2^* allele as a strong candidate for *ATR2^Cala2^*.

### *A2C3^Cala2^* triggers defence in Col-0

When luciferase assays were performed to evaluate the recognition of A2C3^Cala2^ in *Arabidopsis*, a reduction of at least 5-fold in luciferase activity was detected in Col-0 compared to empty vector (EV) control. Equal luciferase activity was detected in an *Hpa*-susceptible recombinant inbred line *Arabidopsis* CW84 when leaf tissue was bombarded with 35S:*A2C3^Cala2^* or EV. These results suggest that Col-0 but not CW84 can recognize A2C3^Cala2^. As before, CCG28 was recognized by WRR4A which served as a positive control (Fig. 2c) (Redkar et al., 2023). Thus, our genetic investigations and bombardment experiments are consistent with A2C3^Cala2^ being the avirulence determinant ATR2^Cala2^, and we hence refer to A2C3^Cala2^ as ATR2^Cala2^. As an additional test of ATR2^Cala2^ detection by RPP2 in *Arabidopsis*, *ATR2*^Cala2^ under 35S promoter was transformed into Col-0. Only three T_1_ lines were selected from antibiotic screening, and strikingly all three transformants showed strong dwarf phenotype, consistent with recognition of *ATR2^Cala2^* in *Arabidopsis* Col-0 background (Fig. 2d).

### *ATR2^Cala2^* enhances susceptibility in the absence of host recognition

Plant pathogen effector proteins that are translocated into host cells can attenuate host defence. Many pathogen effectors interfere with cellular processes that are essential for innate immunity.

To evaluate its virulence function, *ATR2^Cala2^* was transiently expressed in *N. benthamiana* leaves that were then inoculated with *P. infestans* race 88069. The *P. infestans* lesion area was significantly larger in *ATR2^Cala2^*-expressing leaf sectors than in GFP vector control (Fig. 3a). At 7 dpi, lesion area in the *ATR2^Cala2^*-expressing region was more than 4 times larger than that observed in GFP control region (Fig. 3b). Stable *ATR2^Cala2^*-expressing *Arabidopsis* lines (35S:*ATR2^Cala2^*) were generated in Ler-0, which lacks *RPP2A* and *RPP2B*. In contrast to *ATR2^Cala2^* expressing Col-0, all transgenic lines selected grew similar to Ler-0 wild-type (Fig. 3c). Strikingly, all the transgenic lines were more susceptible to virulent *Pst* DC3000 or *Hpa*-Cala2 compared to Ler-0 wild-type control (Fig. 3d, e). Ler-*eds1* was used as hypersusceptible control. Collectively, these data show that in both *Arabidopsis* and *N. benthamiana*, *ATR2^Cala2^* expression can compromise plant innate immunity in the absence of recognition by a cognate *R*-gene.

**Figure 3.**
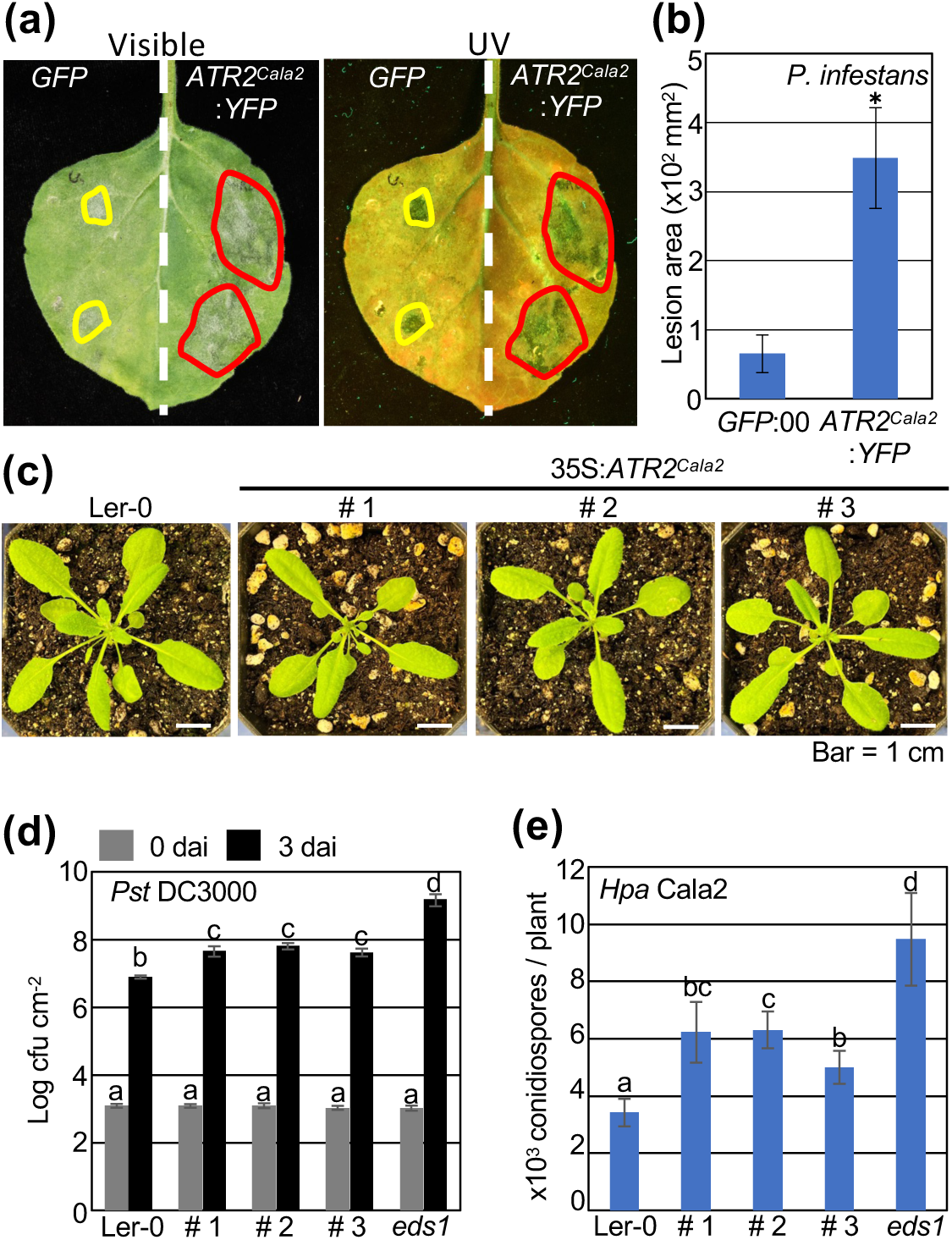
Enhanced disease susceptibility resulting from exogenous *ATR2^Cala2^* expression. (a) Phenotypes on *N. benthamiana* transiently expressing the *GFP* control (*GFP*) or *ATR2^Cala2^*:*YFP* under the 35S promotor followed by *P. infestans* 88069 inoculation (2 x 10^4^ zoospores / ml) 2 days after transient expression. Photos were taken 7 dai with *P. infestans*. (b) Disease lesion area by *P. infestans* on *N. benthamiana* leaves. 40 lesion squares of each were measured. (c) Generation of constitutively *ATR2^Cala2^* expressing transgenic *Arabidopsis* in Ler-0 background. Bar = 1 cm. (d) Bacterial growth in Ler-0, *ATR2^Cala2^*-OX Ler-0 (# 1, # 2 and # 3) and Ws-*eds1* (*eds1*) as a hyper-susceptible control infected with *Pst* DC3000 (10^5^ cfu / ml). (e) Quantification of conidiospores on wild-type and transgenic plants at 7 dai infected with *Hpa* Cala2 (5 x 10^4^ conidiospores / ml). Data are means ± standard deviations from three independent experiments. Asterisks indicate significant differences as determined by Student’s t-test (P < 0.05). According to Fisher’s Least Significant Difference, LSD (P < 0.05), statistical significance was shown by different letters above each bar.

### In addition to RPP2A and RPP2B, two additional linked TNLs, RPP2C and RPP2D, are required for full RPP2 function

We tested the requirement for *RPP2A* (*At1g19500*) and *RPP2B* (*At1g19510*) in ATR2^Cala2^ recognition. There are two other adjacent TIR-NB-LRR genes (*At1g19520* and *At4g19530*, hereafter *RPP2C* and *RPP2D*) (Fig. S8a) that comprise a gene pair similar to *RRS1* and *RPS4*, with C-terminal extended post-LRR domains and a head-to-head orientation (Fig. S8a, b). RPP2A contains two TIR-NB-ARC domains connected by the *Arabidopsis* LSH1 and *Oryza* G1 (ALOG) domain followed by LRRs towards its C-terminus. Post-LRR (PL) domains of RPP2B and RPP2D are homologous to the RPP1 C-terminal jelly-roll/Ig-like (C-JID) domain. RPP2C harbours an additional TIR domain following an extended post-LRR domain (Fig. S8b; Table S4). We obtained the fast neutron 2 (FN2) *rpp2a* mutant (Sinapidou et al, 2004), and several T-DNA insertional mutants from GABI or SALK for *rpp2b*, *rpp2c* and *rpp2d* (Fig. S8c). We combined sequence capture with Illumina sequencing (RenSeq) with DNA from the FN2 (*rpp2a-1*) mutant and confirmed a 25 bp deletion in *RPP2A*. The *RPP2B*, *RPP2C* and *RPP2D* mutations were also verified (Fig. S9). After inoculating mutants with *Hpa*-Cala2, conidiospores were counted at 7 dpi. Ler-0 and Ws-*eds1* were used as susceptible controls. While fewer than 1x10^2^ spores/plant were detected in the resistant Col-0, around 4x10^3^ spores/plant were detected on *rpp2a-1* and *rpp2b-1* mutants with similar values to those obtained from Ler-0 (near 4.8x10^3^/plant), indicating Cala2 resistance in Col-0 is compromised by *rpp2a* or *rpp2b* mutations (Fig. 4a). Interestingly, *rpp2c-1* and *rpp2d-1* mutants also showed compromised resistance to *Hpa*-Cala2. Around 1.2 × 10^3^ to 1.3 × 10^3^ spores/plant were counted from *rpp2c* and *rpp2d* mutants, suggesting *RPP2C* and *RPP2D* also contribute to full resistance against *Hpa*-Cala2 in Col-0 (Fig. 4a). Trailing necrosis was observed on *rpp2c* and *rpp2d* mutants, while no necrosis was observed on Col-0 at 6 dpi (Fig. S10a). To visualize cell death and hyphal growth, we performed trypan blue staining at 5 dpi using infected cotyledons. Local cell death was observed on Col-0, and hyphal growth and haustoria formation over the whole leaf was observed on *rpp2a*, *rpp2b* and Ler-0 cotyledons, as well as Ws-*eds1*. Partial but restricted hyphal growth was detected on *rpp2c* and *rpp2d* mutants (Fig. 4b). When *Hpa*-Emoy2 was inoculated on to *rpp2a*, *rpp2b*, *rpp2c* and *rpp2d* mutants, resistance was not compromised, due to *RPP4*-dependent resistance in Col-0 (van der Biezen et al., 2002; Fig. S11), which is why the susceptible phenotypes are specific to *Hpa*-Cala2.

**Figure 4.**
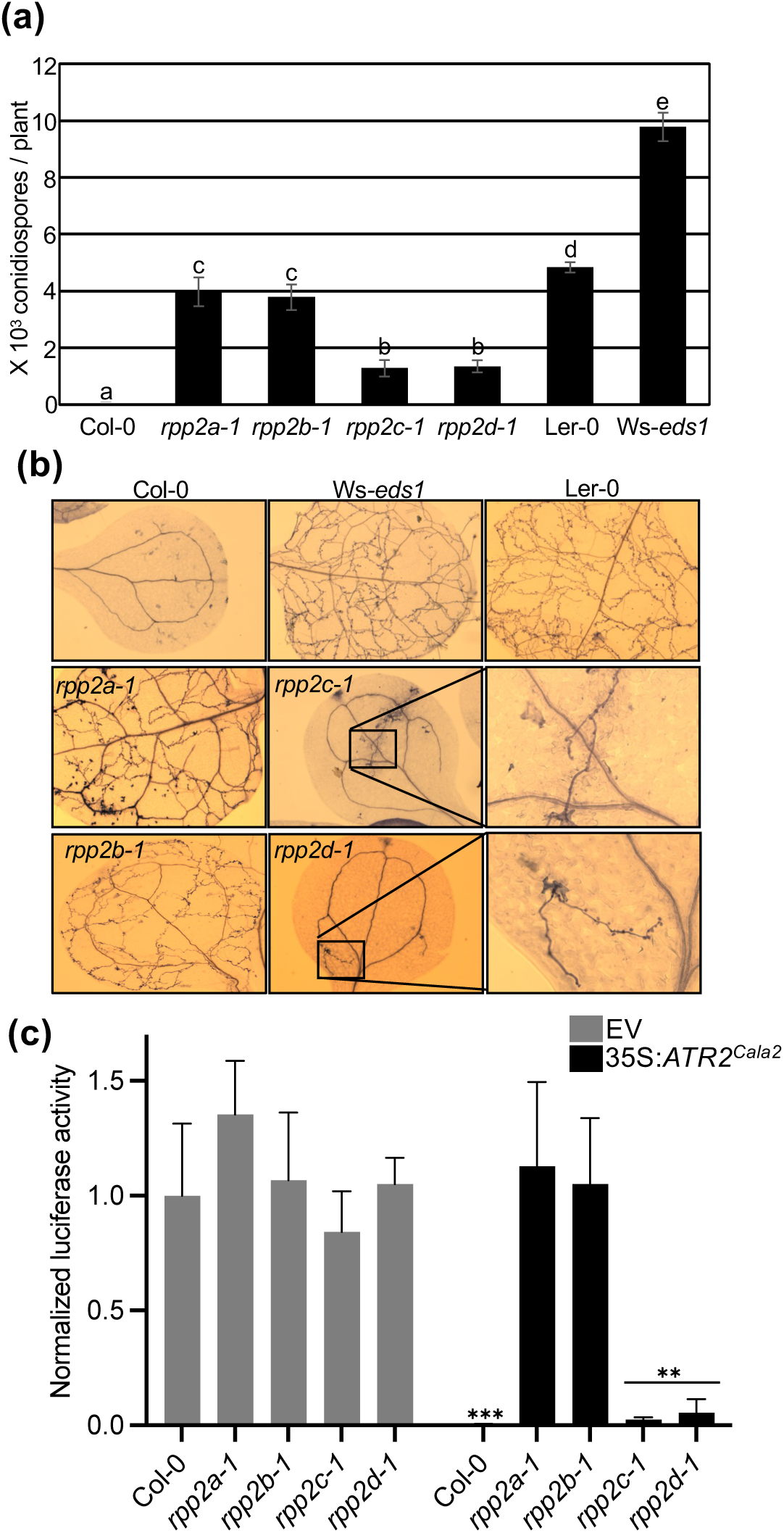
Compromised *Hpa* Cala2 resistance in *rpp2* mutants. (a) Quantification of conidiospores on Col-0, individual *rpp2* mutants from Col-0, Ler-0 and Ws-*eds1* at 7 dai infected with *Hpa* Cala2 (5 x 10^4^ conidiospores / ml). Data are means ± standard deviations from three independent experiments. According to Fisher’s Least Significant Difference, LSD (P < 0.05), statistical significance was shown by different letters above each bar. (b) Trypan blue staining of *Hpa* hyphal growth on cotyledons at 5 dai. Hyphal growth region on *rpp2c* and *rpp2d* mutants was enlarged to clearly show the *Hpa* hyphal development. (c) Luciferase measurement upon biolistic bombardment into Col-0 and *rpp2* mutants. Statistical significance compared with luciferase alone in Col-0 is indicated by asterisks (**, P < 0.01; ***, P < 0.001) according to two-way ANOVA with Tukey’s multiple comparison test.

To assess *ATR2^Cala2^* recognition capacity by RPP2 paralogs, luciferase eclipse assays were conducted using individual Col-0 *rpp2a-1*, *rpp2b-1*, *rpp2c-1* and *rpp2d-1* mutants. The luciferase activity was normalized to compare with that of EV control on Col-0 (Fig. 4c). The normalized luciferase activity in each individual Col-0 *rpp2a* mutant and Col-0 with EV was comparable with no significant differences indicating particle bombardment distributed well through the leaves of Col-0 and each mutant. When *ATR2^Cala2^* was bombarded together with 35S:*luciferase* on Col-0, normalized luciferase activity was strongly reduced, ranging from 0.004 to 0.007, while those on *rpp2a-1* or *rpp2b-1* still maintained a range of 0.86 to 1.55, indicating ATR2^Cala2^ recognition is almost completely abolished in *rpp2a-1* and *rpp2b-1* mutants. ATR2^Cala2^ was still recognized in *rpp2c-1* and *rpp2d-1* mutants, with normalized activity ranging from 0.01 to 0.1 (Fig. 4c). Even though no statistically significant differences were detected between *ATR2^Cala2^*-bombarded Col-0, *rpp2c-1* and *rpp2d-1*, the average values of the luciferase activities on *rpp2c-1* (mean, 0.024) and *rpp2d-1* (mean, 0.054) are almost 5-10 times higher than on Col-0 (mean, 0.005) when co-bombarded with *ATR2^Cala2^*, consistent with *RPP2C* and *RPP2D* weakly contributing to ATR2^Cala2^ recognition.

Transgenic complementation assays with CW84 were carried out using JAtY 49E17 clone (Zhou et al., 2011), which harbours the whole RPP2 cluster. While sporangiophore formation was observed on CW84, complemented transgenic plants restored complete resistance to *Hpa*-Cala2 (Fig. S10b).

We transiently expressed *ATR2^Cala2^* in *N. benthamiana* with *RPP2A* and/or *RPP2B*, but no hypersensitive response (HR) was observed at 3 dpi (Fig. S12a). HR-like cell death was observed when *ATR2^Cala2^* allele was transiently co-expressed with *RPP2A*, *RPP2B*, *RPP2C* and *RPP2D* at 5 dpi (Fig. S12b). These data indicate *RPP2C* and *RPP2D* are also required for full ATR2^Cala2^-triggered immunity.

### RPP2 haplotype diversity

As the *RPP2* cluster containing *RPP2A*, *RPP2B*, *RPP2C* and *RPP2D* is required for full ATR2^Cala2^-triggered resistance, we assessed *RPP2* haplotype diversity in multiple *Arabidopsis* accessions. An investigation of the *Arabidopsis* pan-NLRome (Van de Weyer et al., 2019) enabled in-depth analysis for the *RPP2* cluster. We compared the *RPP2* cluster in 64 *A. thaliana* accessions (Fig. S13). Col-0, Oy-0, and Can-0 have the complete form of the *RPP2* cluster, while other accessions lack some *RPP* genes or harbor incomplete (partial) *RPP* genes (Fig. 5a). Interestingly, almost all accessions contain a complete form of *RPP2B*, and 7 ecotypes among 21 harbor *RPP2A* while many of other ecotypes have incomplete alleles of *RPP2A* (Fig. 5a). Almost half of accessions lack, or contain partial forms of, *RPP2C* or *RPP2D* (Fig. 5a). Among 64 accessions, while only 17 accessions contain complete *RPP2A* encoding TIR-NB-TIR-NB-LRR (TN-TNL) homologs, the *RPP2B*-encoding TIR-NB-LRR (TNL) is well conserved in almost all accessions excluding Ler-0, Rsch-4 and Vig-1 (Fig. S13, S14a). *RPP2C* is lacking or incomplete in more than 40 accessions and the amino acid length of RPP2D is quite diverse (Fig. S13). While RPP2A haplotypes show structural diversity on their first TIR-NB-ARC domains, RPP2B, RPP2C and RPP2D are more conserved in different *Arabidopsis* accessions (Fig. S14).

**Figure 5.**
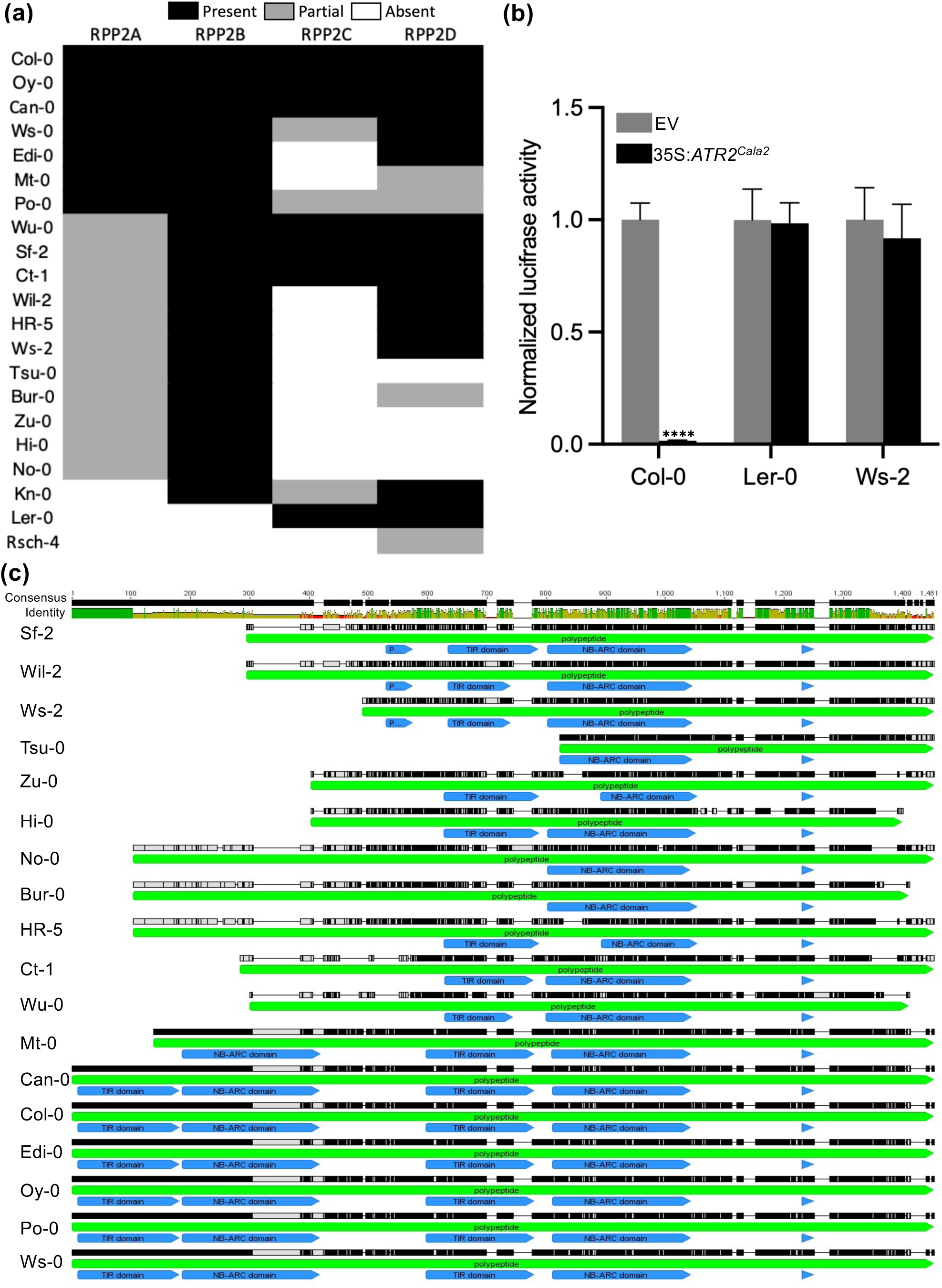
Differential *ATR2^Cala2^* recognition capacity dependent on RPP2A haplotype. (a) Heatmap diagram for *RPP2* cluster haplotype analyses from 21 *Arabidopsis* accessions. (b) Normalized luciferase activity by biolistic bombardment of *ATR2^Cala2^* with luciferase into Col-0, Ler-0 (*RPP2A, 2B*-lacking) and Ws-2 (partial *RPP2A*). Data are means ± standard deviations from three independent experiments. Asterisk indicates a significant difference as determined by two-way ANOVA with Tukey’s test (****, P < 0.0001). (c) RPP2A haplotype analyses from 21 Arabidopsis ecotypes.

To monitor ATR2^Cala2^ recognition capacity, luciferase eclipse assay bombardment was conducted with Col-0 (complete RPP2 cluster), Ler-0 (*RPP2C* and *RPP2D* lacking), and Ws-2 (partial *RPP2A* and *RPP2C* lacking). As expected, ATR2^Cala2^ is recognized in Col-0, while Ler-0 and Ws-2 lack ATR2^Cala2^ recognition capacity indicating a critical role of *RPP2A* and *RPP2B* for ATR2^Cala2^ recognition (Fig. 5c). Compared to other *RPP2* genes, *RPP2A* was present in diverse forms. As shown in Fig. 5c, diverse accessions including Ws-2 lost the first N-terminal TIR-NB-ARC domain. The TIR-NB-ARC defect in Ws-2 abolishes ATR2^Cala2^ recognition.

## Discussion

Downy mildews such as *B. lactucae* on lettuce (Parra et al., 2021), *P. viticola* on grapevines (Li et al., 2015), *P. cubensis* on cucumber (Zhang et al., 2019) and *H. brassicae* on brassicas (Liu et al., 2021) are destructive obligate oomycete phytopathogens on fruit and vegetable crops (Thines and Kamoun, 2010; Tör et al., 2023). Genetic variation for downy mildew resistance has also been studied in *Brassica* species such as *B. napus*, broccoli, non-heading Chinese cabbage, and Chinese cabbage (Chen et al., 2008; Xiao et al., 2016). The 27 known *Dm* genes in lettuce are located in gene clusters that encode NLRs (Parra et al., 2021). A better understanding of resistance mechanisms to downy mildew is highly desirable. *Arabidopsis* NLR-encoding *RPP* genes confer recognition of specific downy mildew races and different RPP proteins specifically recognize their cognate downy mildew RxLR effectors (Asai et al., 2018).

Most oomycete pathogens deploy secreted effector proteins, with the signature amino acid motif RxLR, which enter plant cells where they promote virulence (Win et al., 2012; Asai et al., 2014; Wood et al., 2020). The function and evolution of RxLR effectors have been investigated since their discovery (Anderson et al., 2015). Comparative genomics indicates that *RxLR* genes play a major role in virulence for downy mildews and *Phytophthora* species. Although progress has been made, there is still much to learn about the mechanisms of downy mildew virulence and host resistance. Most *P. infestans* and *Hpa* effectors carry an RxLR motif.

We positionally identified *ATR2^Cala2^* that encodes a non-canonical RxLR-like protein recognized by RPP2A and RPP2B. *ATR2^Cala2^*encodes an RxLR-like protein with an N-terminal signal peptide, and dEER and C-terminal Y and WY modules. The *ATR2* alleles in other *Hpa* strains are identical to the *ATR2^Emoy2^* allele and lack GHVR, dEER and WY motifs due to a frame shift resulting from a single nucleotide deletion. In the absence of recognition by *RPP2A* and *RPP2B*, *ATR2^Cala2^* expression enhances pathogen susceptibility *in planta*. Furthermore, the head-to-head *RPP2C* and *RPP2D* genes which are adjacent to *RPP2A* and *RPP2B* also contribute to full resistance to *Hpa*-Cala2. ATR5 was the first example of a non-canonical RxLR effector lacking the canonical RxLR motif but with an N-terminal signal peptide and a canonical EER motif (Bailey et al., 2011). At the expected RxLR position, ATR5 carries Gly-Arg-Val-Arg (GRVR) instead of RxLR. ATR2^Cala2^ at this position carries Gly-His-Val-Arg (GHVR) followed by a dEER motif. ATR5 contains two WY motifs and one LWY motif at its C-terminus (Fig. S15a). The Y-WY domain of ATR2^Cala2^ resembles the LWY of ATR5 based on Alphafold2 structural prediction (Fig. S15b). In the *Hpa* genome, more than 150 genes encode for potentially secreted proteins like ATR2^Cala2^ that carry motifs such as signal peptide and EER but lack the RxLR motif (Asai et al., 2014). This also been seen in other oomycetes such as *<u>Pseudoperonospora</u>* and *Bremia* (Purayannur et al., 2020; Wood et al., 2020; Nur et al., 2023). We conclude that although the RxLR motif is often found in oomycete effectors, in *Hpa* as in other oomycetes, some divergence is permitted for effector translocation. Therefore, additional *Hpa* effectors may exist that have not yet been predicted.

*RPP2* was the first genetically defined *R*-gene locus shown to carry two NLR-encoding genes, both of which are required for function (Sinapidou et al., 2004). The corresponding recognized effector from *Hpa* enables investigations into how the *RPP2*-encoded immune receptor complex functions. Recognized effectors are also valuable tools for investigating plant/microbe interactions, since their host targets correspond to important plant defense components. Most *R* gene pair-encoding NLR proteins, such as *Arabidopsis* RRS1-RPS4 that recognize bacterial effectors AvrRps4 and PopP2, and rice RGA4-RGA5 recognizing rice blast effectors AVR1-CO39 and AVR-Pia, are encoded by divergently transcribed genes (Cesari et al., 2013; Ma et al., 2018), in contrast to *RPP2A* and *RPP2B* (Fig. S8a). Sensor NLRs are dependent on executor (or helper) NLRs for downstream immune signalling (Feehan et al., 2020). RRS1 functions as a sensor that reveals effectors that target WRKY domain transcription factors, while RPS4 is an executor (Ma et al., 2018). Uniquely, RPP2A contains two TIR-NB-ARC domains followed by LRR, and an ALOG domain which is specific and conserved to land plants and has DNA-binding activity (Yoshida et al., 2009; Naramoto et al., 2020; Beretta et al., 2023) between the two TIR-NB-ARC domains (Fig. S8). Comparative analyses of RPP2A in diverse *Arabidopsis* accessions show that the main variation in the RPP2A haplotype is the presence or absence of one N-terminal TIR-NB-ARC (Fig. 5c, S13). RPP2B is a typical TIR-NLR resembling the executor NLR, RPS4. Compared to RPP2A, RPP2B is relatively well conserved in different *Arabidopsis* accessions (Fig. S13, S14a). Conceivably, RPP2A functions as a sensor for ATR2^Cala2^ and RPP2B functions as a signal executor. TIR domains of plant NLRs are known to have nicotinamide adenine dinucleotide hydrolase (NADase) activity, which requires a catalytic glutamate (E), that activates defense (Wan et al., 2019). The C-JID domains of *Arabidopsis* RPP1 and *N. benthamiana* ROQ1 (recognition of XopQ1) are required for pathogen effector recognition. ATR1 binds to the C-JID and the LRRs of RPP1 leading to assembly of tetramers with NADase activity (Ma et al., 2020). The LRR and C-JID of ROQ1 directly interact with *Xanthomonas* effector (XopQ) allowing the NB-ARC domain to transition to an ATP-bound state. Complex assembly results in TIR proximity that opens the NADase active site (Martin et al., 2020). The first TIR on RPP2A has the catalytic E residue whereas the second TIR on RPP2A lacks the conserved E residue (Table S4). The TIR of RPP2B has the conserved E residue and a C-JID (Table S4), consistent with a role as executor. Still, the domains of RPP2A and/or RPP2B that interact with ATR2^Cala2^ remain to be elucidated.

We also revealed the requirement for two additional *TIR-NB-LRR* genes, *RPP2C* and *RPP2D*, adjacent to *RPP2A* and *RPP2B* and showed all 4 NLR proteins are required for full resistance against *Hpa*-Cala2. A paired head-to-head *R*-gene structure is often found in plant paired NLRs (Narusaka et al., 2009; Cesari et al., 2014; Saucet et al., 2021). *RPP2C* and *RPP2D* form a head-to-head orientation similar to *RRS1*-*RPS4* (Narusaka et al., 2009; Ma et al., 2018; Guo et al., 2020). The C-terminal post-LRR domain of RPS4 is homologous with C-JID suggesting that it recognizes conformational changes in RRS1 upon effector recognition (Saucet et al., 2021). The RPP2C post-LRR domain is homologous to that of RRS1 but RPP2C contains a TIR domain on its C-terminal end instead of WRKY. RPP2D contains a C-JID on its C-terminus homologous to that of RPS4 and RPP1 (Fig. S8; Table S4). Many *Arabidopsis* accessions lack or have incomplete RPP2C but RPP2D is relatively conserved among different accessions (Fig. S13). Even though *RPP2C* and *RPP2D* are quantitatively required for full resistance against *Hpa*-Cala2, how this pair contributes to ATR2^Cala2^ recognition remains unclear. Conceivably, their contribution could either be additive, or by potentiating RPP2A/B-dependent recognition. We speculate that since both RPP2A and RPP2C carry integrated TIR domains which do not contain a catalytic E residue, ATR2^Cala2^ might function by interacting with and somehow suppressing functions of host TIR domain-containing proteins. Interestingly, the TIR on RPP2C C-terminal end lacks the catalytic E residue for NADase activity, while the first TIR on RPP2C has the E residue, and RPP2D also has the E residue on its TIR (Table S4). We hypothesize that the second TIR on RPP2A might function to detect ATR2^Cala2^ leading to conformational change via RPP2B interaction with ATR2^Cala2^. This could result in RPP2A/B resistosome activation enabling signal transduction through activated NADase function of the first TIR on RPP2A and RPP2B TIR. If the TIR in RPP2A lacking catalytic E functions as an effector decoy, the C-terminal TIR on RPP2C might also act as an integrated decoy to detect ATR2^Cala2^. However, thus far we were unable to detect direct or indirect interaction between ATR2^Cala2^ and each RPP2 protein. Further research is needed to define the effector recognition mechanisms for these atypical NLR protein pairs.

## Acknowledgement

Financial support from the Gatsby Foundation (http://www.gatsby.org.uk/), and from BBSRC grants BB/K009176/1 and BB/M003809/1 to JDG Jones, is gratefully acknowledged. This work is also supported in parts by the grant 09 963/A from the Leverhulme Trust to M. Tör. We thank Matthew Smoker and Jodie Taylor for their help with *Arabidopsis* transformation. The authors would like to thank Dr. Kenichi Tsuda for providing luciferase assay kit for DSK.

## Author contributions

DSK, MT and JDGJ conceptualized and designed the research. DSK, AW-T, VC and OJF conducted all experiments. DSK and MT performed the data analysis. VC and MT gave critical intellectual input and provided materials for this work. YL and WM carried out structural prediction and analyses of *Hpa* effectors. DSK, MT and JDGJ wrote the manuscript with input from all co-authors.

## Data availability

All the sequence data used in this study can be found in NCBI (See Materials and Methods). The data supporting the findings of the study are available from the corresponding author upon request.

## References

1. Allen RL, Bittner-Eddy PD, Grenville-Briggs LJ, Meitz JC, Rehmany AP, Rose LE, Beynon JL. 2004. Host-parasite coevolutionary conflict between *Arabidopsis* and downy mildew. Science 306(5703): 1957–1960.

2. Anderson RG, Deb D, Fedkenheuer K, McDowell JM. 2015. Recent progress in RXLR effector research. Molecular Plant-Microbe Interactions 28(10): 1063–1072.

3. Asai S, Furzer OJ, Cevik V, Kim DS, Ishaque N, Goritschnig S, Staskawicz BJ, Shirasu K, Jones JDG. 2018. A downy mildew effector evades recognition by polymorphism of expression and subcellular localization. Nature Communications 9(1): 5192.

4. Asai S, Rallapalli G, Piquerez SJM, Caillaud MC, Furzer OJ, Ishaque N, Wirthmueller L, Fabro G, Shirasu K, Jones JDG. 2014. Expression profiling during Arabidopsis/downy mildew interaction reveals a highly-expressed effector that attenuates responses to salicylic acid. PLoS Pathogens 10(10): e1004443.

5. Bailey K, Cevik V, Holton N, Byrne-Richardson J, Sohn KH, Coates M, Woods-Tör A, Aksoy HM, Hughes L, Baxter L, et al. 2011. Molecular cloning of ATR5^Emoy2^ from *Hyaloperonospora arabidopsidis*, an avirulence determinant that triggers RPP5-mediated defense in *Arabidopsis*. Molecular Plant-Microbe Interactions 24(7): 827–838.

6. Baxter L, Tripathy S, Ishaque N, Boot N, Cabral A, Kemen E, Thines M, Ah-Fong A, Anderson R, Badejoko W, et al. 2010. Signatures of adaptation to the obligate biotrophy in the *Hyaloperonospora arabidopsidis* genome. Science 330(6010): 1549–1551.

7. Bernoux M, Ve T, Willams S, Warren C, Hatters D, Valkov E, Zhang X, Ellis JG, Kobe B, Dodds PN. 2011. Structural and functional analysis of a plant resistance protein TIR domain reveals interfaces for self-association, signaling, and autoregulation. Cell Host & Microbe 9(3): 200–211.

8. Birch PRJ, Rehmany AP, Pritchard L, Kamoun S, Beynon JL. 2006. Trafficking arms: oomycete effectors enter host plant cells. Trends in Microbiology 14(1): 8–11.

9. Bomblies K, Lempe J, Warthmann N, Lanz C, Dangl JL, Weigel D. 2007. Autoimmune response as a mechanism for a Dobzhansky-Muller-type incompatibility syndrome in plants. PLoS Biology 5(9): e236.

10. Botella MA, Parker JE, Frost LN, Bittner-Eddy PD, Beynon JL, Daniels MJ, Holub EB, Jones JDG. 1998. Three genes of the Arabidopsis *RPP1* complex resistance locus recognize distinct *Peronospora parasitica* avirulence determinants. Plant Cell 10(11): 1847–1860.

11. Boutrot F, Zipfel C. 2017. Function, discovery, and exploitation of plant pattern recognition receptors for broad-spectrum disease resistance. Annual Review of Phytopathology 55: 257–286.

12. Césari S, Kanzaki H, Fujiwara T, Bernoux M, Chalvon V, Kawano Y, Shimamoto K, Dodds P, Terauchi R, Kroj T. 2014. The NB-LRR proteins RGA4 and RGA5 interact functionally and physically to confer disease resistance. EMBO Journal 33(17): 1941–1959.

13. Césari S, Thilliez G, Ribot C, Chalvon V, Michel C, Jauneau A, Rivas S, Alaux L, Kanzaki H, Okuyama Y, Morel JB, Fournier E, Tharreau D, Terauchi R, Kroj T. 2013. The rice resistance protein pair RGA4/RGA5 recognizes the *Magnaporthe oryzae* effectors AVR-Pia and AVR1-CO39 by direct binding. Plant Cell 25(4):1463–1481.

14. Chen XF, Hou XL, Zhang JY, Zheng JQ. 2008. Molecular characterization of two important antifungal proteins isolated by downy mildew infection in non-heading Chinese cabbage. Molecular Biology Reports 35(4): 621–629.

15. Chisholm ST, Coaker G, Day B, Staskawicz BJ. 2006. Host-microbe interactions: shaping the evolution of the plant immune response. Cell 124(4): 803–814.

16. Clough SJ, Bent AF. 1998. Floral dip: a simplified method for *Agrobacterium*-mediated transformation of *Arabidopsis thaliana*. The Plant Journal 16(6): 735–743.

17. Coates ME, Beynon JL. 2010. *Hyaloperonospora arabidopsidis* as a pathogen model. Annual Review of Phytopathology 48: 329–345.

18. Dangl JL, Horvath DM, Staskawicz BJ. 2013. Pivoting the plant immune system from dissection to deployment. Science 341(6147): 746–751.

19. Dodds PN, Rathjen JP. 2010. Plant immunity: towards an integrated view of plant-pathogen interactions. Nature Reviews Genetics 11(8): 539–548.

20. Eitas TK, Dangl JL. 2010. NB-LRR proteins: pairs, pieces, perceptions, partners, and pathways. Current Opinion in Plant Biology 13(4): 472–477.

21. Feehan JM, Castel B, Bentham AR, Jones JDG. 2020. Plant NLRs get by with a little help from their friends. Current Opinion in Plant Biology 56: 99–108.

22. Feng F, Zhou JM. 2012. Plant-bacterial pathogen interactions mediated by type III effectors. Current Opinion in Plant Biology 15(4): 469–476.

23. Geu-Flores F, Nour-Eldin HH, Nielsen MT, Halkier BA. 2007. USER Fusion: A rapid and efficient method for simultaneous fusion and cloning of multiple PCR products. Nucleic Acids Research 35(7): e55.

24. Goritschnig S, Krasileva KV, Dahlbeck D, Staskawicz BJ. 2012. Computational prediction and molecular characterization of an oomycete effector and the cognate *Arabidopsis* resistance gene. PLoS Genetics 8(2): e1002502.

25. Guo H, Ahn HK, Sklenar J, Huang J, Ma Y, Ding P, Menke FLH, Jones JDG. 2020. Phosphorylation-regulated activation of the *Arabidopsis* RRS1-R/RPS4 immune receptor complex reveals two distinct effector recognition mechanisms. Cell Host & Microbe 27(5): 769–781.

26. He J, Ye W, Choi DS, Wu B., Zhai Y, Guo B., Duan S., Wang Y, Gan J, Ma W, Ma J. 2019. Structural analysis of *Phytophthora* suppressor of RNA silencing 2 (PSR2) reveals a conserved modular fold contributing to virulence. *Proceedings of the National Academy of Sciences*, USA 116(16): 8054–8059

27. Høie MH, Kiehl EN, Petersen B, Nielsen, M, Winther O, Nielsen H, Hallgren J, Marcatili P. 2022. NetSurfP-3.0: accurate and fast prediction of protein structural features by protein language models and deep learning. Nucleic Acids Research 50(W1): W510–W515.

28. Holub EB. 2008. Natural history of *Arabidopsis thaliana* and oomycete symbioses. European Journal of Plant Pathology 122: 91–109.

29. Holub EB, Beynon JL, Crute IR. 1994. Phenotypic and genotypic characterisation of interactions between isolates of *Peronospora parasitica* and accessions of *Arabidopsis thaliana*. Molecular Plant-Microbe Interactions 7(2): 223–239.

30. Hou Y, Zhai Y, Feng L, Karimi HZ, Rutter BD, Zeng L, Choi DS, Zhang B, Gu W, Chen X, et al. 2019. A *Phytophthora* effector suppresses trans-kingdom RNAi to promote disease susceptibility. Cell Host & Microbe 25(1):153–165

31. Jones DA, Takemoto D. 2004. Plant innate immunity -direct and indirect recognition of general and specific pathogen-associated molecules. Current Opinion in Immunology 16(1): 48–62.

32. Jones JDG, Dangl JL. 2006. The plant immune system. Nature 444(7117): 323–329.

33. Jones JDG, Vance RE, Dangl JL. 2016. Intracellular innate immune surveillance devices in plants and animals. Science 354(6316): aaf6395

34. Jumper J, Evans R, Pritzel A, Green T, Figurnov M, Ronneberger O, Tunyasuvunakool K, Bates R, Žídek A, Potapenko A, et al. 2021. Highly accurate protein structure prediction with AlphaFold. Nature 596(7873): 583–589.

35. Kim DS, Kim NH, Hwang BK. 2015. Glycine-rich RNA-binding protein1 interacts with receptor-like cytoplasmic protein kinase1 and suppresses cell death and defense responses in pepper (*Capsicum annuum*). New Phytologist 205(2): 786–800.

36. Kover PX, Valdar W, Trakalo J, Scarcelli N, Ehrenreich IM, Purugganan MD, Durrant C, Mott R. 2009. A multiparent advanced generation inter-cross to fine-map quantitative traits in *Arabidopsis thaliana*. PLoS Genetics 5(7): e1000551.

37. Krasileva KV, Dahlbeck D, Staskawicz BJ. 2010. Activation of an *Arabidopsis* resistance protein is specified by the in planta association of its leucine-rich repeat domain with the cognate oomycete effector. Plant Cell 22(7): 2444–2458.

38. Li H, Yu SC, Zhang FL, Yu YJ, Zhao XY, Zhang DS, Zhao X. 2011. Development of molecular markers linked to the resistant QTL for downy mildew in *Brassica Rapa* L. Ssp. Pekinensis. Hereditas 33(11): 1271–1278.

39. Li X, Wu J, Yin L, Zhang YL, Qu JJ, Lu J. 2015. Comparative transcriptome analysis reveals defense-related genes and pathways against downy mildew in *Vitis amurensis* grapevine. Plant Physiology and Biochemistry 95: 1–14.

40. Liu Y, Li D, Yang N, Zhu X, Han K, Gu R, Bai J, Wang A, Zhang Y. 2021. Genome-side identification and analysis of CC-NBS-LRR family in response to downy mildew and black rot in Chinese cabbage. International Journal of Molecular Sciences 22(8): 4266.

41. Ma S, Lapin D, Liu L, Sun Y, Song W, Zhang X, Logemann E, Yu D, Wang J, Jirschitzka J, et al. 2020. Direct pathogen-induced assembly of an NLR immune receptor complex to form a holoenzyme. Science 370(6521): eabe3069.

42. Ma Y, Guo H, Hu L, Martinez PP, Moschou PN, Cevik V, Ding P, Duxbury Z, Sarris PF, Jones JDG. 2018. Distinct modes of derepression of an Arabidopsis immune receptor complex by two different bacterial effectors. Proceedings of the National Academy of Sciences, USA 115(41): 10218–10227.

43. Martin R, Qi T, Zhang H, Liu F, King M, Toth C, Staskawicz BJ. 2020. Structure of the activated Roq1 resistosome directly recognizing the pathogen effector XopQ. Science 370(6521): eabd9993.

44. Meyers BC, Kozik A, Kuang H, Michelmore RW. 2003. Genome-wide analysis of NBS-LRR-encoding genes in Arabidopsis. Plant Cell 15(4): 809–834.

45. Monaghan J, Zipfel C. 2012. Plant pattern recognition receptor complexes at the plasma membrane. Current Opinion in Plant Biology 15(4): 349–357.

46. Naramoto S, Hata Y, Kyozuka J. 2020. The origin and evolution of the ALOG proteins, members of a plant-specific transcription factor family, in land plants. Journal of Plant Research 133(3): 323–329.

47. Narusaka M, Shirasu K, Noutoshi Y, Kubo Y, Shiraishi T, Iwabuchi M, Narusaka Y. 2009. RRS1 and RPS4 provide a dual Resistance-gene system against fungal and bacterial pathogens. The Plant Journal 60(2): 218–226.

48. Nur MJ, Wood KJ, Michelmore RW. 2023. EffectorO: motif-independent prediction of effectors in oomycete genomes using machine learning and lineage-specificity. Molecular Plant-Microbe Interactions doi: 10.1094/MPMI-11-22-0236-TA.

49. Nürnberger T, Brunner F, Kemmerling B, Piater L. 2004. Innate immunity in plants and animals: striking similarities and obvious differences. Immunolocal Reviews 198: 249–266.

50. Oliver RP, Ipcho SV. 2004. Arabidopsis pathology breathes new life into the necrothrphs- vs.-biotrophs classification of fungal pathogens. Molecular Plant Pathology 5(4): 347–352.

51. Parra L, Nortman K, Sah A, Truco MJ, Ochoa O, Michelmore R. 2021. Identification and mapping of new genes for resistance to downy mildew in lettuce. Theoretical and Applied Genetics 134(2): 519–528.

52. Punta M, Coggill PC, Eberhardt RY, Mistry J, Tate J, Boursnell C, Pang N, Forslund K, Ceric G, Clements J, et al. 2012. The Pfam protein families database. Nucleic Acids Research 40(D1): D290–D301.

53. Purayannur S, Cano LM, Bowman MJ, Childs KL, Gent DH, Quesada-Ocampo LM. 2020. The effector repertoire of the hop downy mildew pathogen *Pseudoperonospora humuli*. Frontiers in Genetics 11:910.

54. Redkar A, Cevik V, Bailey K, Zhao H, Kim DS, Zou Z, Furzer OJ, Fairhead S, Borhan MH, Holub EB, Jones JDG. 2023. The Arabidopsis WRR4A and WRR4B paralogous NLR proteins both confer recognition of multiple *Albugo candida* effectors. New Phytologist 237(2): 532–547.

55. Rehmany AP, Gordon A, Rose LE, Allen RL, Armstrong MR, Whisson SC, Kamoun S, Tyler BM, Birch PR, Beynon JL. 2005. Differential recognition of highly divergent downy mildew avirulence gene alleles by *RPP1* resistance from two Arabidopsis lines. Plant Cell 17(6): 1839–1850

56. Rehmany AP, Grenville LJ, Gunn ND, Allen RL, Paniwnyk Z, Byrne J, Whisson SC, Birch PR, Beynon JL. 2003. A genetic interval and physical contig spanning the *Peronospora parasitica* (At) avirulence gene locus ATR1Nd. Fungal Genetic Biology 38(1): 33–42.

57. Saucet SB, Esmenjaud D, Van Ghelder C. 2021. Integrity of the post-LRR domain is required for TIR-NB-LRR function. Molecular Plant-Microbe Interactions 34(3): 286–296.

58. Sarris PF, Duxbury Z, Huh SU, Ma Y, Segonzac C, Sklenar J, Derbyshire P, Cevik V, Rallapalli G, Saucet SB, et al. 2015. A plant immune receptor detects pathogen effectors that target WRKY transcription factors. Cell 161(5): 1089–1100.

59. Sinapidou E, Williams K, Nott L, Bahkt S, Tör M., Crute I, Bittner-Eddy P, Beynon J. 2004. Two TIR:NB:LRR genes are required to specify resistance to *Peronospora parasitica* isolate Cala2 in *Arabidopsis*. The Plant Journal 38(6): 898–909.

60. Slusarenko AJ, Schlaich NL. 2003. Downy mildew of *Arabidopsis thaliana* caused by *Hyaloperonospora parasitica* (formerly *Peronospora parasitica*). Molecular Plant Pathology 4(3): 159–170.

61. Sohn KH, Lei R, Nemri A, Jones JDG. 2007. The downy mildew effector proteins ATR1 and ATR13 promote disease susceptibility in *Arabidopsis thaliana*. Plant Cell 19(12): 4077–4090.

62. Spoel SH, Dong X. 2012. How do plants achieve immunity? Defence without specialized immune cells. Nature Review Immunology 12(2): 89–100.

63. Thines M, Kamoun S. 2010. Oomycete–plant coevolution: recent advances and future prospects. Current Opinion in Plant Biology 13(4): 427–433.

64. Tör M, Wood T, Webb A, Göl D, McDowell JM. 2023. Recent developments in plant-downy mildew interactions. Seminars in Cell and Developmental Biology doi: 10.1016/j.semcdb.2023.01.010.

65. Van de Weyer AL, Monteiro F, Furzer OJ, Nishimura MT, Cevik V, Witek K, Jones JDG, Dangl JL, Weigel D, Bemm F. 2019. A species-wide inventory of NLR genes and alleles in *Arabidopsis thaliana*. Cell 178(5): 1260–1272.

66. van der Biezen EA, Freddie CT, Kahn K, Parker JE, Jones JDG. 2002. *Arabidopsis RPP4* is a member of the *RPP5* multigene family of TIR-NB-LRR genes and confers downy mildew resistance through multiple signalling components. The Plant Journal 29(4): 439–451.

67. Varadi V, Anyango S, Deshpande M, Nair S, Natassia C, Yordanova G, Yuan D, Stroe O, Wood G, Laydon A, et al. 2021. AlphaFold Protein Structure Database: massively expanding the structural coverage of protein-sequence space with high-accuracy models. Nucleic Acids Research 50(D1): D439–D444.

68. Wan L, Essuman K, Anderson RG, Sasaki Y, Monteiro R, Chung EH, Nishimura EO, DiAntonio A, Milbrandt J, Dangl JL, et al. 2019. TIR domains of plant immune receptors are NAD^+^-cleaving enzymes that promote cell death. Science 365(6455): 799–803.

69. Win J, Krasileva KV, Kamoun S, Shirasu K, Staskawicz BJ, Banfield MJ. 2012. Sequence divergent RXLR effectors share a structural fold conserved across plant pathogenic oomycete species. PLoS Pathogens 8(1): e1002400.

70. Win J, Morgan W, Bos J, Krasileva KV, Cano LM, Chaparro-Garcia A, Ammar R, Staskawicz BJ, Kamoun S. 2007. Adaptive evolution has targeted the C-terminal domain of the RXLR effectors of plant pathogenic oomycetes. Plant Cell 19(8): 2349–2369.

71. Wood KJ, Nur M, Gil J, Fletcher K., Lakeman K, Gann D, Gothberg A, Khuu T, Kopetzky J, Naqvi S, et al. 2020. Effector prediction and characterization in the oomycete pathogen *Bremia lactucae* reveal host-recognized WY domain protein that lack the canonical RXLR motif. PLoS Pathogens 16(10); e1009012.

72. Woods-Tör A, Studholme DJ, Cevik V, Telli O, Holub EB, Tör M. 2018. A suppressor/avirulence gene combination in *Hyaloperonospora arabidopsidis* determines race specificity in *Arabidopsis thaliana*. Frontiers in Plant Science 9: 265.

73. Xiao D, Liu ST, Wei YP, Zhou DY, Hou XL, Li Y, Hu, CM. 2016. cDNA-AFLP analysis reveals differential gene expression in incompatible interaction between infected non-heading Chinese cabbage and *Hyaloperonospora parasitica*. Horticulture Research 3: 16034.

74. Xiong A, Ye W, Choi DS, Wong J, Qiao Y, Tao K, Wang Y, Ma W. 2014. Phytophthora suppressor of RNA silencing 2 is a conserved RxLR effector that promotes infection in soybean and *Arabidopsis thaliana*. Molecular Plant-Microbe Interactions 27(12): 1379–1389.

75. Yoshida A, Suzaki T, Tanaka W, Hirano HY. 2009. The homeotic gene *long sterile lemma* (*G1*) specifies sterile lemma identity in the rice spikelet. Proceedings of the National Academy of Sciences, USA 106(47): 20103–20108.

76. Zhang P, Zhu Y, Luo X, Zhou S. 2019. Comparative proteomic analysis provides insights into the complex responses to *Pseudoperonospora cubensis* infection of cucumber (*Cucumis sativus* L.). Scientific Reports 9(1): 9433.

77. Zhou R, Benavente LM, Stepanova AN, Alonso JM. 2011. A recombineering-based gene tagging system for Arabidopsis. The Plant Journal 66(4): 712–723.

