## Supporting information for "ATR2^Cala2^ from *Arabidopsis*-infecting downy mildew requires 4 TIR-NLR immune receptors for full recognition"

### *New Phytologist* Supporting Information

The following Supporting Information is available for this article:

**Fig. S1** Map-based cloning approach for *ATR2^Cala2^* allele.

**Fig. S2** Diagram of a contig in which *ATR2* candidates co-segregate.

**Fig. S3** Loci, structure and recognition of *A2C1* and *A2C2*.

**Fig. S4** *A2C1* and *A2C2* do not trigger cell death with *RPP2* in *Nicotiana* species*.*

**Fig. S5** Determination of *A2C3^Cala2^* allele cosegregation on avirulent *Hpa* Cala2-Noks1 F2 population.

**Fig. S6** Protein sequence analyses of A2C3^Cala2^.

**Fig. S7** *A2C3* expression after *Hpa* Emoy2 and Cala2 infection.

**Fig. S8** Scheme of RPP2 cluster.

**Fig. S9** RenSeq-MiSeq from *rpp2a* mutant.

**Fig. S10** Compromised ATR2-mediated resistance in *rpp2* mutants.

**Fig. S11** *Hpa* Emoy2 growth on Col-0, *rpp2* mutants and Oy-0.

**Fig. S12** HR triggered after transient expression of *ATR2* with *RPP2A*, *2B*, *2C* and *2D* in *N. benthamiana.*

**Fig. S13** RPP2 cluster haplotype analyses from pan-NLRome (Van de Weyer et al., 2019).

**Fig. S14** Haplotype analyses of RPP2B, RPP2C and RPP2D from several *Arabidopsis* accessions. **Fig. S15** Comparison of WY domains of ATR2^Cala2^ and ATR5.

**Table S1** Interaction phenotypes (IP) with CaNo F2 isolates.

**Table S2** Mapping table of *ATR2*.

**Table S3** The number of SNPs on *A2C3* alleles on 7 different *Hpa* isolates.

**Table S4** Domains on RPP2 proteins and catalytic E residue conservation on each TIR.

[Note: if your file is a large table, e.g. an Excel file, this should be submitted separately]

**Methods S1** Supplemental Materials and Methods
