## Supplementary material for "ATR2^Cala2^ from *Arabidopsis*-infecting downy mildew requires 4 TIR-NLR immune receptors for full recognition": Figs_supple

### Slide 1
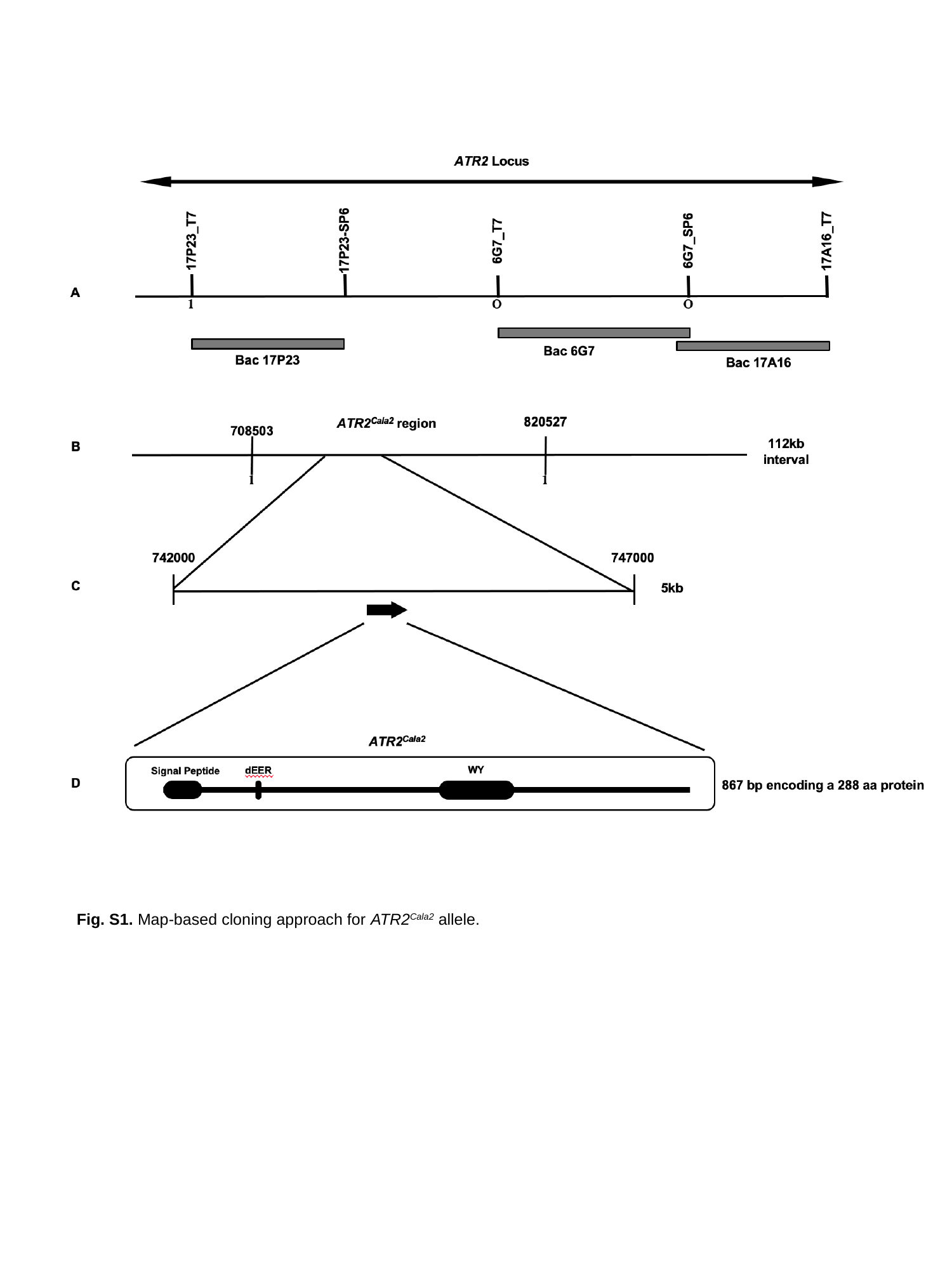

Fig. S1. Map-based cloning approach for ATR2^Cala2^ allele.

### Slide 2
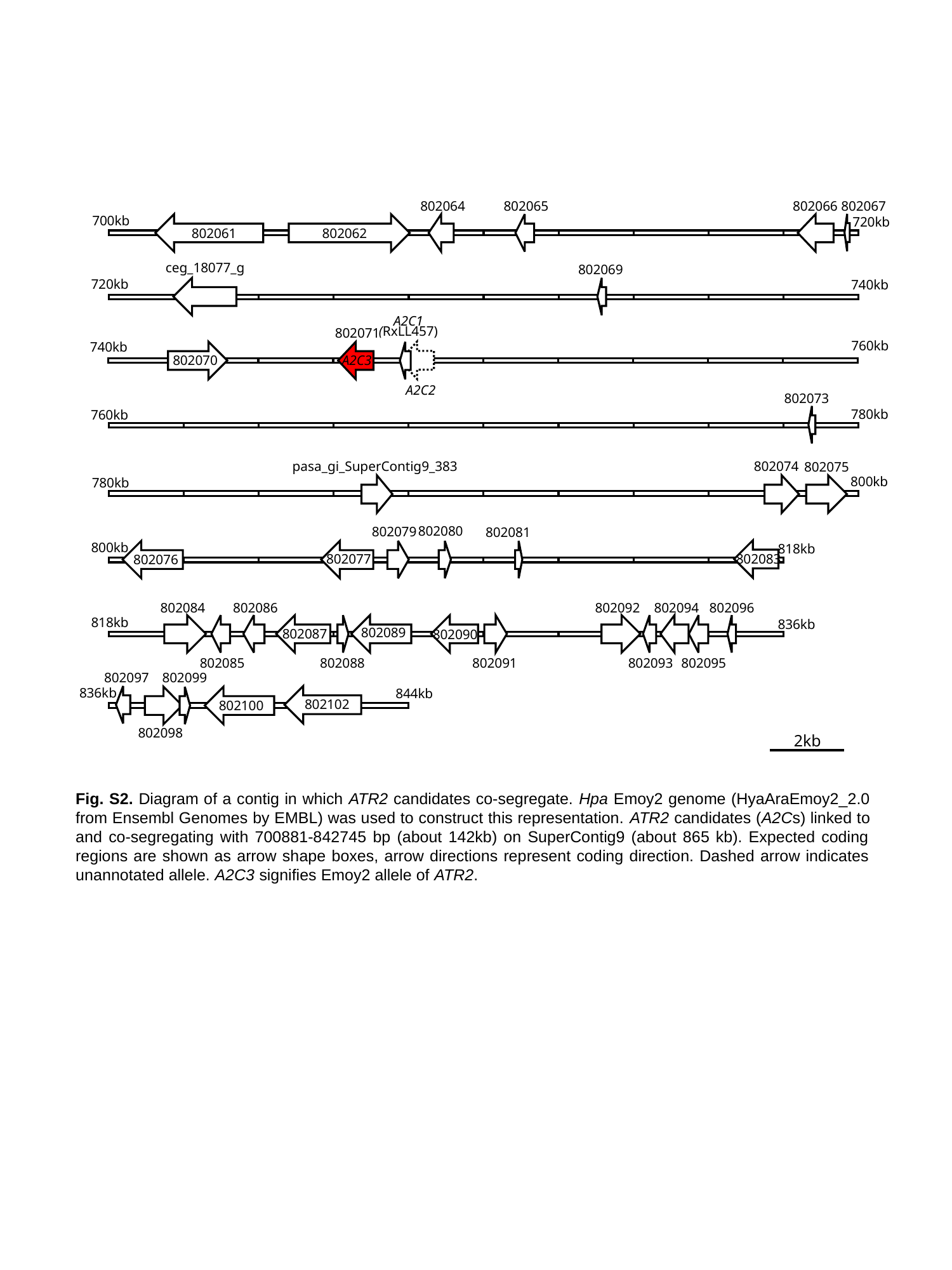

802064
802065
802066
802067
700kb
720kb
802061
802062
ceg_18077_g
802069
720kb
740kb
A2C1
(RxLL457)
802071
760kb
740kb
802070
A2C3
A2C2
802073
780kb
760kb
pasa_gi_SuperContig9_383
802074
802075
800kb
780kb
802080
802079
802081
800kb
818kb
802077
802083
802076
802084
802086
802092
802094
802096
818kb
836kb
802089
802087
802090
802085
802088
802091
802093
802095
802097
802099
836kb
844kb
802102
802100
802098
2kb
Fig. S2. Diagram of a contig in which ATR2 candidates co-segregate. Hpa Emoy2 genome (HyaAraEmoy2_2.0 from Ensembl Genomes by EMBL) was used to construct this representation. ATR2 candidates (A2Cs) linked to and co-segregating with 700881-842745 bp (about 142kb) on SuperContig9 (about 865 kb). Expected coding regions are shown as arrow shape boxes, arrow directions represent coding direction. Dashed arrow indicates unannotated allele. A2C3 signifies Emoy2 allele of ATR2.

### Slide 3
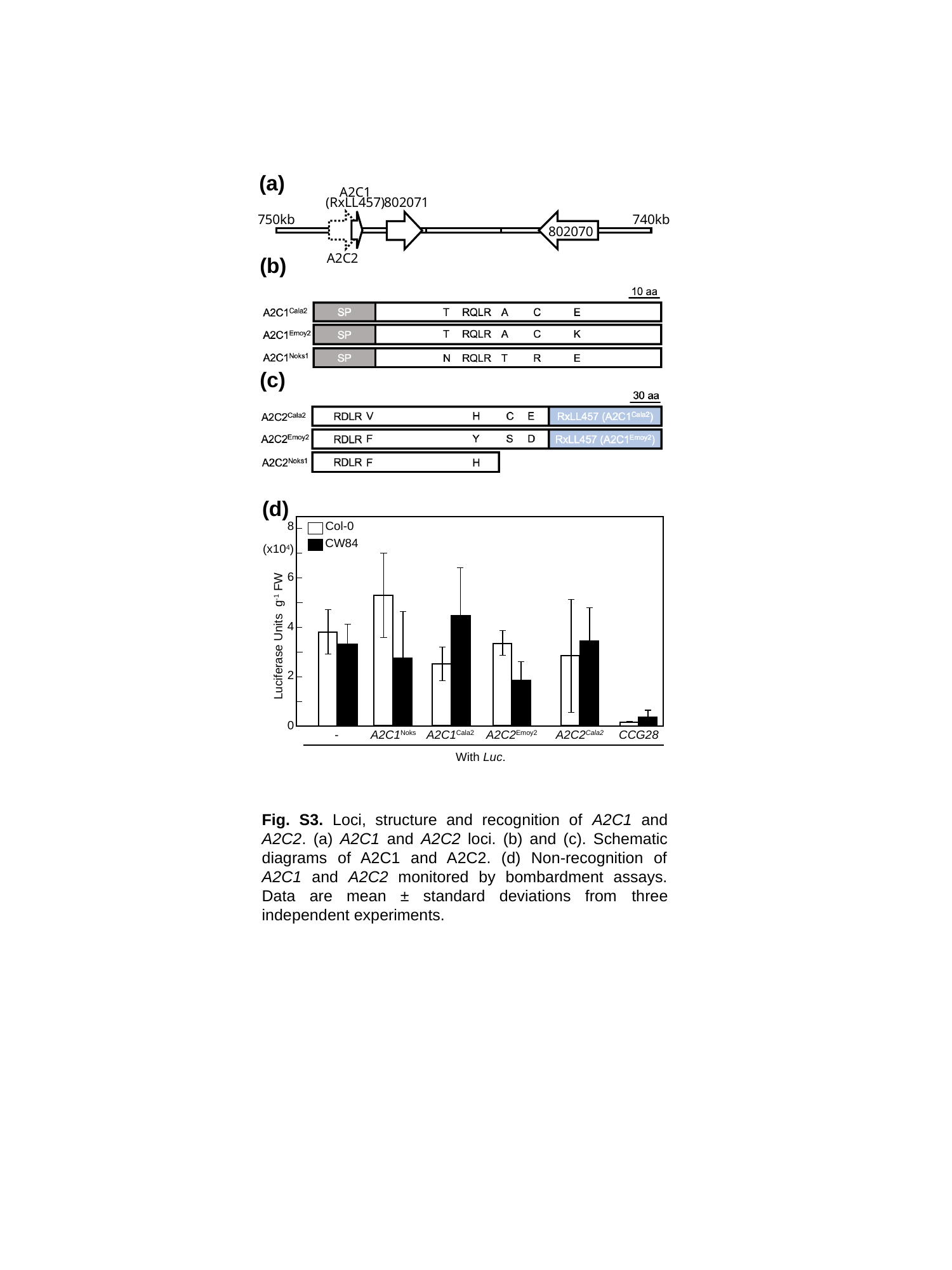

(a)
A2C1
(RxLL457)
802071
750kb
740kb
802070
A2C2
(b)
(c)
(d)
8
Col-0
CW84
(x10^4^)
6
4
Luciferase Units g^-1^ FW
2
0
-
A2C1^Noks^
A2C1^Cala2^
A2C2^Emoy2^
A2C2^Cala2^
CCG28
With Luc.
Fig. S3. Loci, structure and recognition of A2C1 and A2C2. (a) A2C1 and A2C2 loci. (b) and (c). Schematic diagrams of A2C1 and A2C2. (d) Non-recognition of A2C1 and A2C2 monitored by bombardment assays. Data are mean ± standard deviations from three independent experiments.

### Slide 4
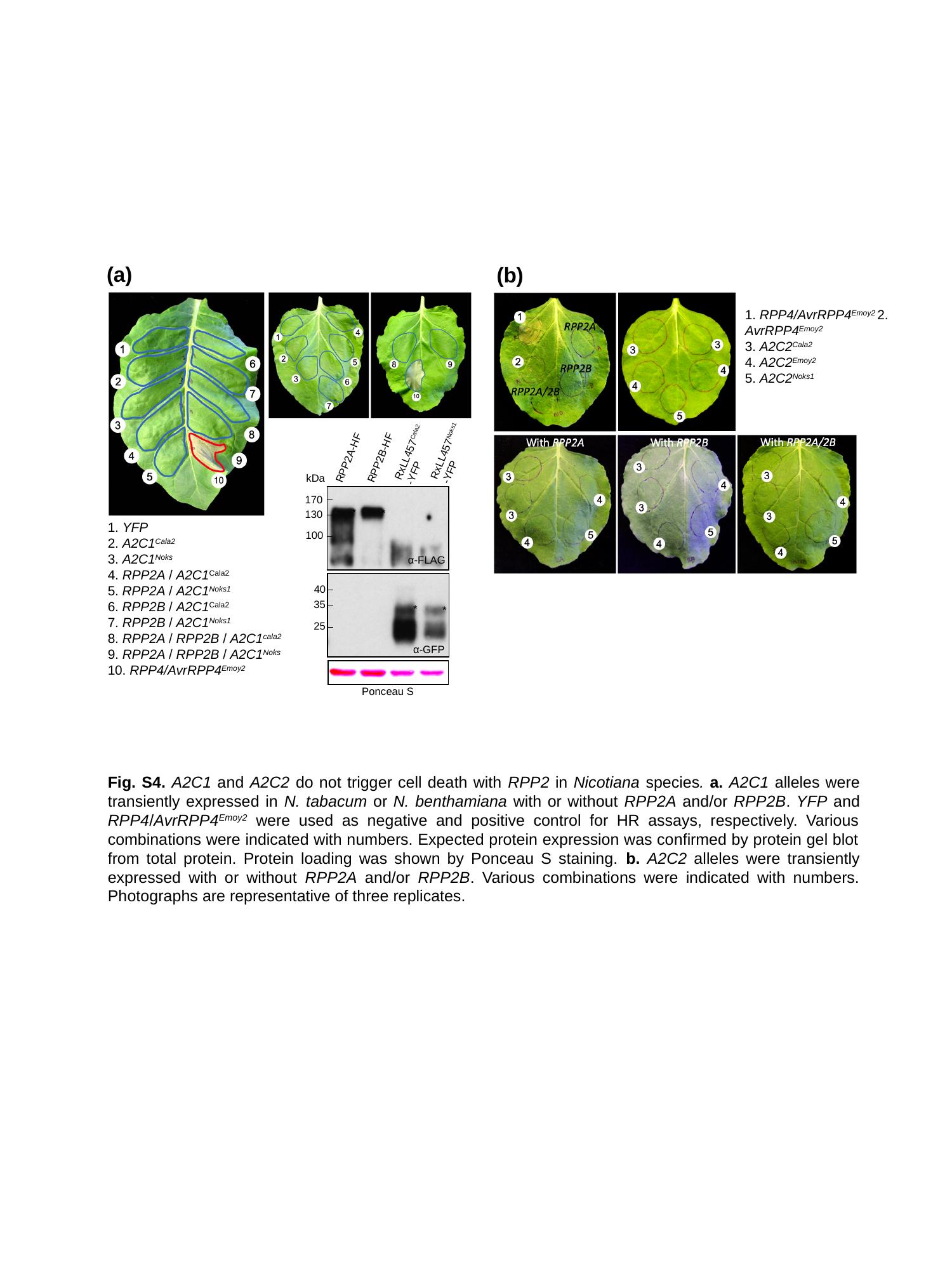

(a)
(b)
1. RPP4/AvrRPP4^Emoy2^ 2. AvrRPP4^Emoy2^
3. A2C2^Cala2^
4. A2C2^Emoy2^
5. A2C2^Noks1^
RxLL457^Noks1^
-YFP
RxLL457^Cala2^
-YFP
RPP2B-HF
RPP2A-HF
kDa
170
*
*
130
100
α-FLAG
40
35
*
*
25
α-GFP
Ponceau S
1. YFP
2. A2C1^Cala2^
3. A2C1^Noks^
4. RPP2A / A2C1^Cala2^
5. RPP2A / A2C1^Noks1^
6. RPP2B / A2C1^Cala2^
7. RPP2B / A2C1^Noks1^
8. RPP2A / RPP2B / A2C1^cala2^
9. RPP2A / RPP2B / A2C1^Noks^
10. RPP4/AvrRPP4^Emoy2^
Fig. S4. A2C1 and A2C2 do not trigger cell death with RPP2 in Nicotiana species. a. A2C1 alleles were transiently expressed in N. tabacum or N. benthamiana with or without RPP2A and/or RPP2B. YFP and RPP4/AvrRPP4^Emoy2^ were used as negative and positive control for HR assays, respectively. Various combinations were indicated with numbers. Expected protein expression was confirmed by protein gel blot from total protein. Protein loading was shown by Ponceau S staining. b. A2C2 alleles were transiently expressed with or without RPP2A and/or RPP2B. Various combinations were indicated with numbers. Photographs are representative of three replicates.

### Slide 5
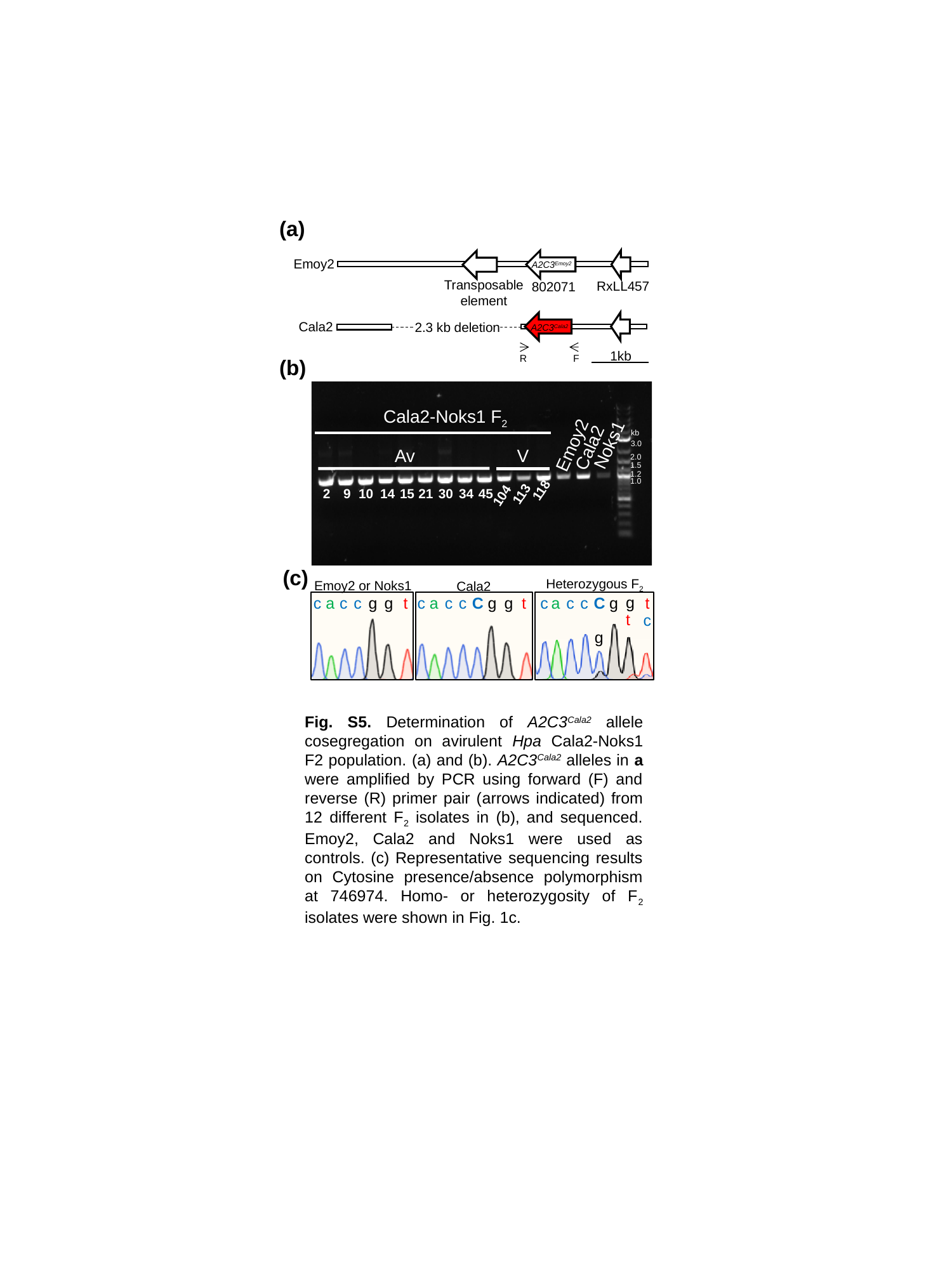

(a)
Emoy2
Transposable
element
RxLL457
802071
A2C3^Emoy2^
Cala2
2.3 kb deletion
A2C3^Cala2^
1kb
R
F
(b)
Cala2-Noks1 F_2_
kb
Noks1
Emoy2
Cala2
3.0
V
Av
2.0
1.5
1.2
1.0
118
2
9
10
14
15
21
30
34
45
113
104
(c)
Heterozygous F_2_
Emoy2 or Noks1
Cala2
c
a
c
c
g
g
t
c
a
c
c
C
g
g
t
g
t
c
a
c
c
C
g
g
t
c
Fig. S5. Determination of A2C3^Cala2^ allele cosegregation on avirulent Hpa Cala2-Noks1 F2 population. (a) and (b). A2C3^Cala2^ alleles in a were amplified by PCR using forward (F) and reverse (R) primer pair (arrows indicated) from 12 different F_2_ isolates in (b), and sequenced. Emoy2, Cala2 and Noks1 were used as controls. (c) Representative sequencing results on Cytosine presence/absence polymorphism at 746974. Homo- or heterozygosity of F_2_ isolates were shown in Fig. 1c.

### Slide 6
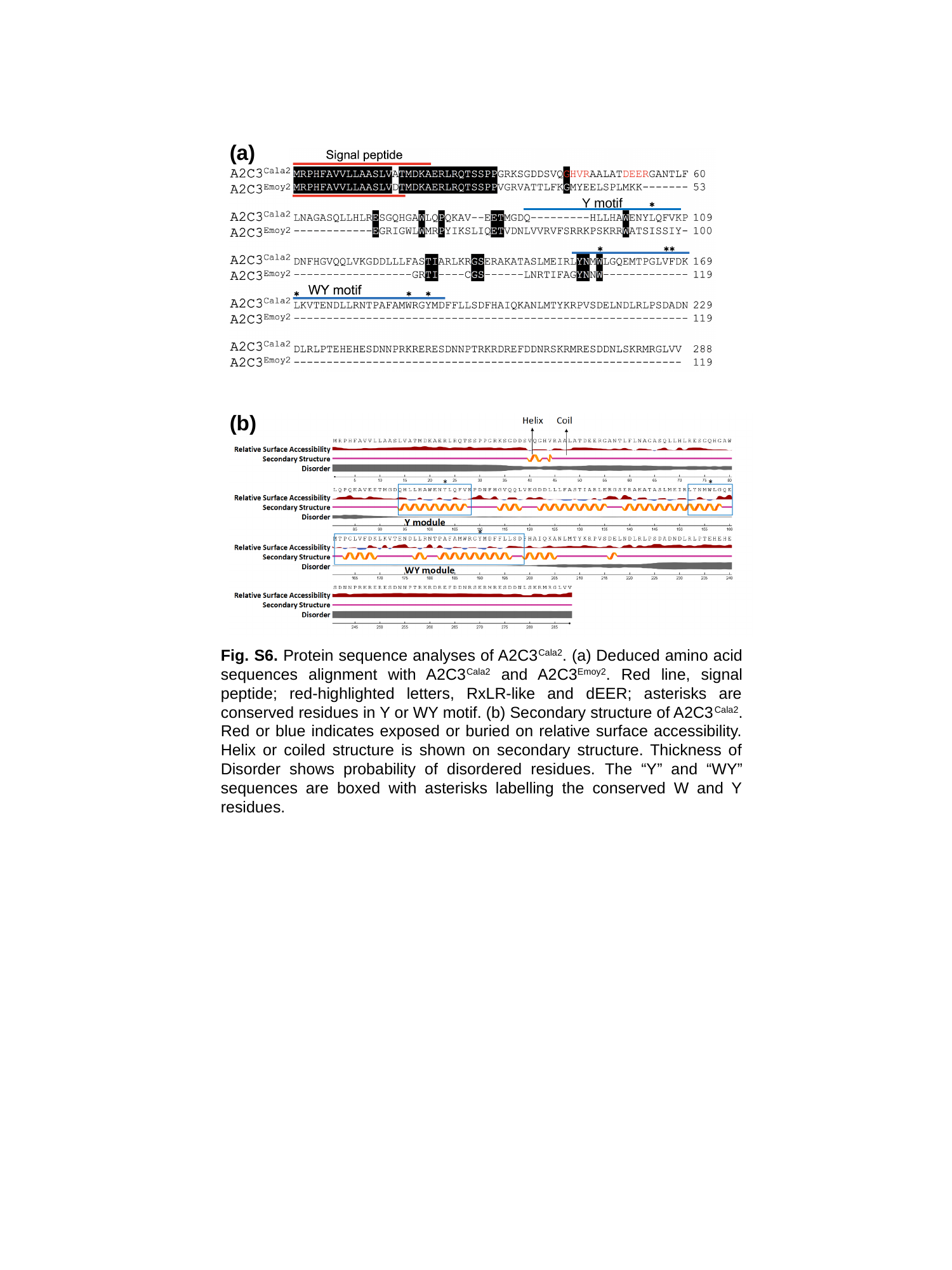

(a)
Y motif
✱
(b)
Fig. S6. Protein sequence analyses of A2C3^Cala2^. (a) Deduced amino acid sequences alignment with A2C3^Cala2^ and A2C3^Emoy2^. Red line, signal peptide; red-highlighted letters, RxLR-like and dEER; asterisks are conserved residues in Y or WY motif. (b) Secondary structure of A2C3^Cala2^. Red or blue indicates exposed or buried on relative surface accessibility. Helix or coiled structure is shown on secondary structure. Thickness of Disorder shows probability of disordered residues. The “Y” and “WY” sequences are boxed with asterisks labelling the conserved W and Y residues.

### Slide 7
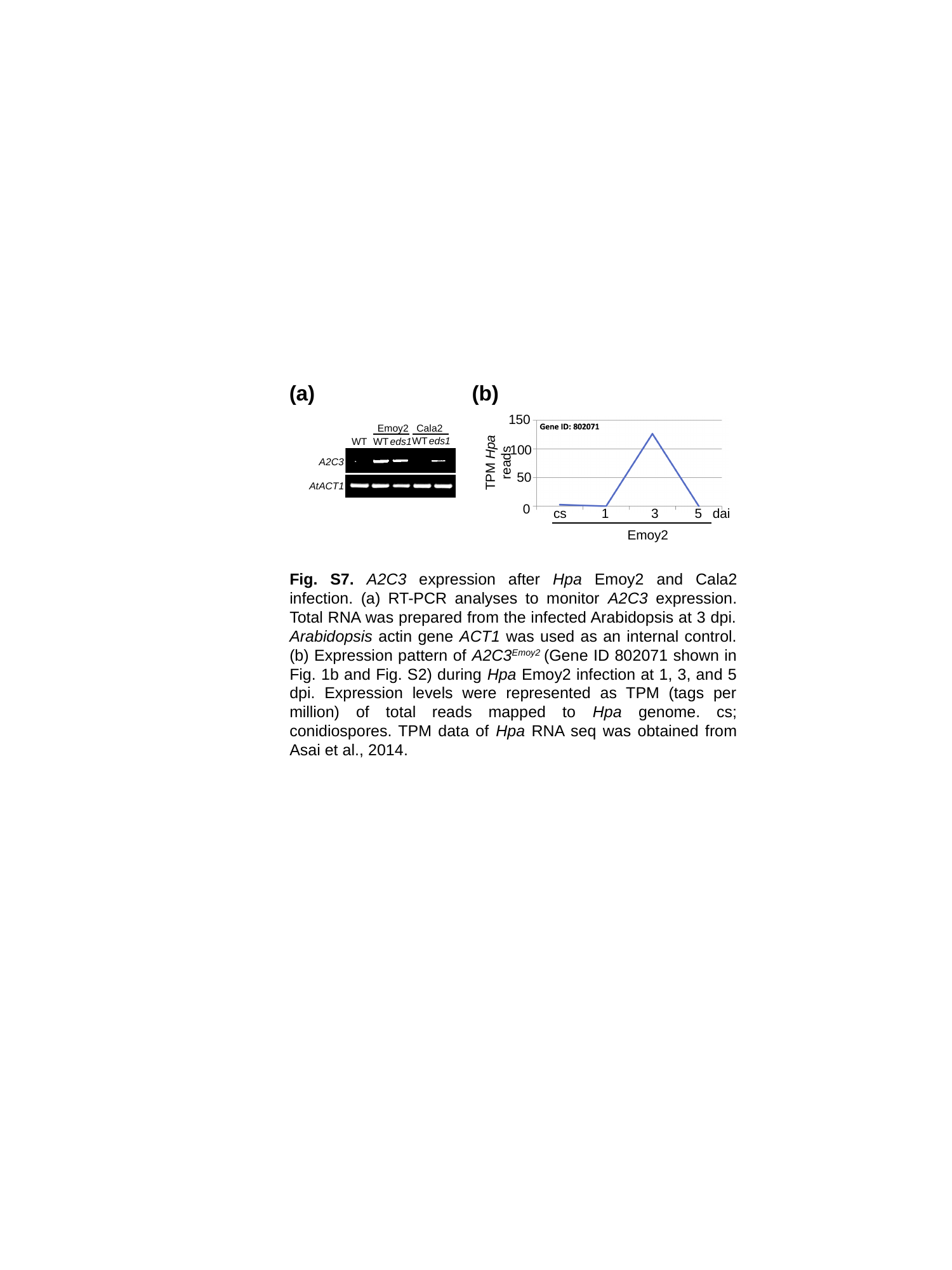

(a)
(b)
150
Emoy2
Cala2
WT
eds1
WT
eds1
WT
100
TPM Hpa reads
A2C3
50
AtACT1
0
cs
1
3
5
dai
Emoy2
Fig. S7. A2C3 expression after Hpa Emoy2 and Cala2 infection. (a) RT-PCR analyses to monitor A2C3 expression. Total RNA was prepared from the infected Arabidopsis at 3 dpi. Arabidopsis actin gene ACT1 was used as an internal control. (b) ﻿Expression pattern of A2C3^Emoy2^ (Gene ID 802071 shown in Fig. 1b and Fig. S2) during Hpa Emoy2 infection at 1, 3, and 5 dpi. Expression levels were represented as TPM (tags per million) of total reads mapped to Hpa genome. cs; conidiospores. TPM data of Hpa RNA seq was obtained from Asai et al., 2014.

### Slide 8
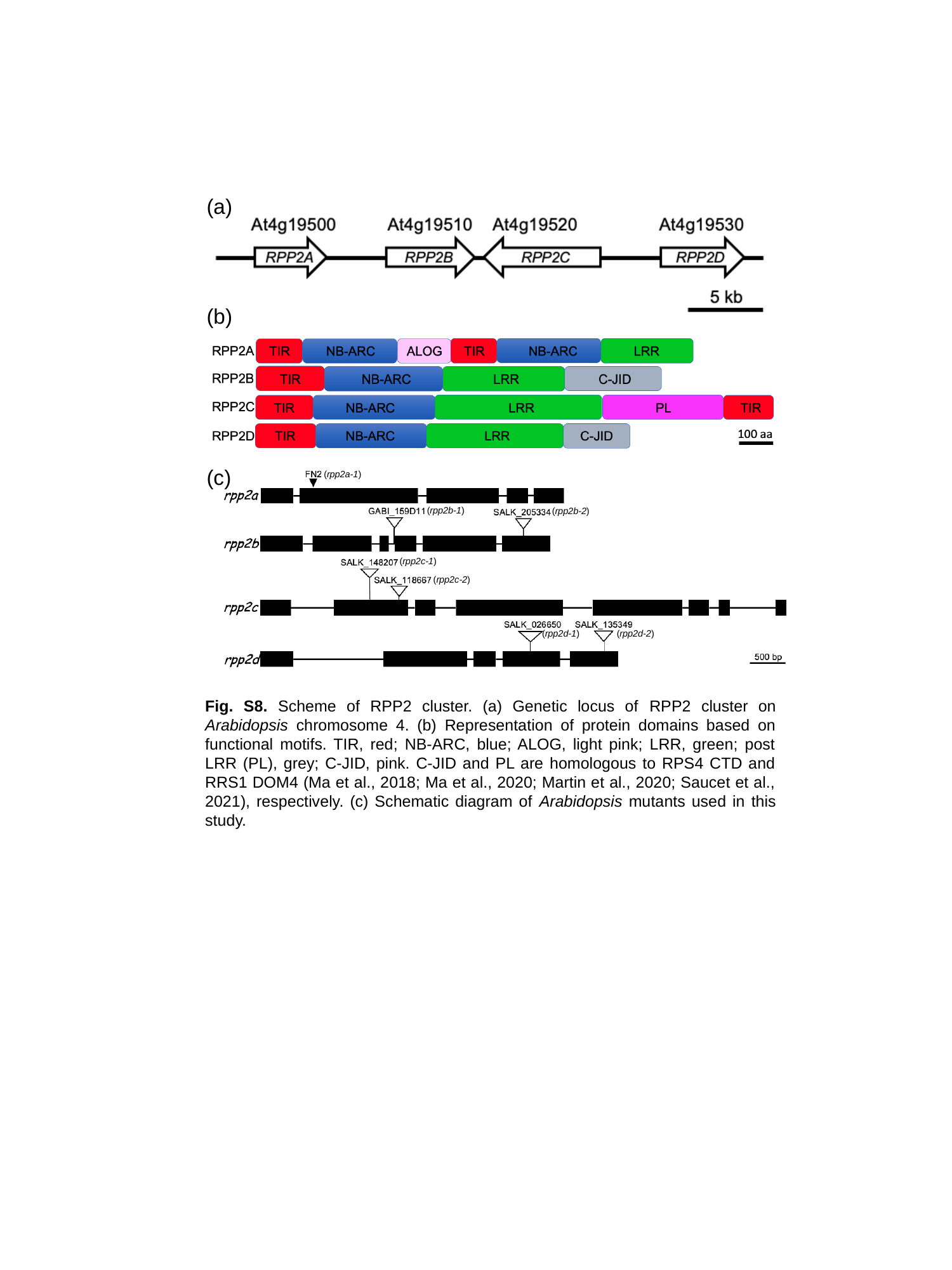

(a)
(b)
(c)
(rpp2a-1)
(rpp2b-1)
(rpp2b-2)
(rpp2c-1)
(rpp2c-2)
(rpp2d-1)
(rpp2d-2)
Fig. S8. Scheme of RPP2 cluster. (a) Genetic locus of ﻿RPP2 cluster on Arabidopsis chromosome 4. (b) ﻿Representation of protein domains based on functional motifs. TIR, red; NB-ARC, blue; ALOG, light pink; LRR, green; post LRR (PL), grey; C-JID, pink. C-JID and PL are homologous to RPS4 CTD and RRS1 DOM4 (Ma et al., 2018; Ma et al., 2020; Martin et al., 2020; Saucet et al., 2021), respectively. (c) Schematic diagram of Arabidopsis mutants used in this study.

### Slide 9
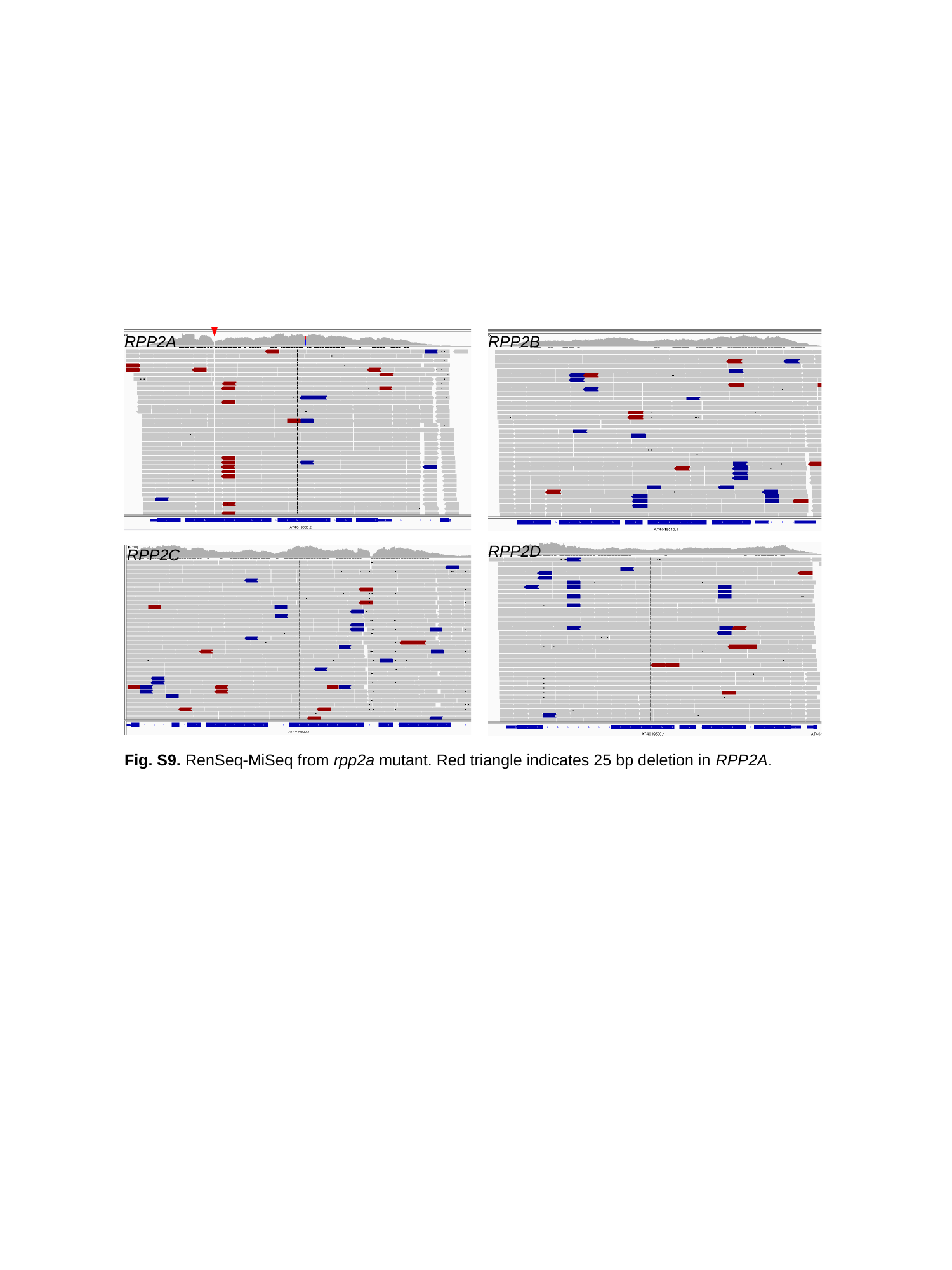

RPP2A
RPP2B
RPP2D
RPP2C
Fig. S9. RenSeq-MiSeq from rpp2a mutant. Red triangle indicates 25 bp deletion in RPP2A.

### Slide 10
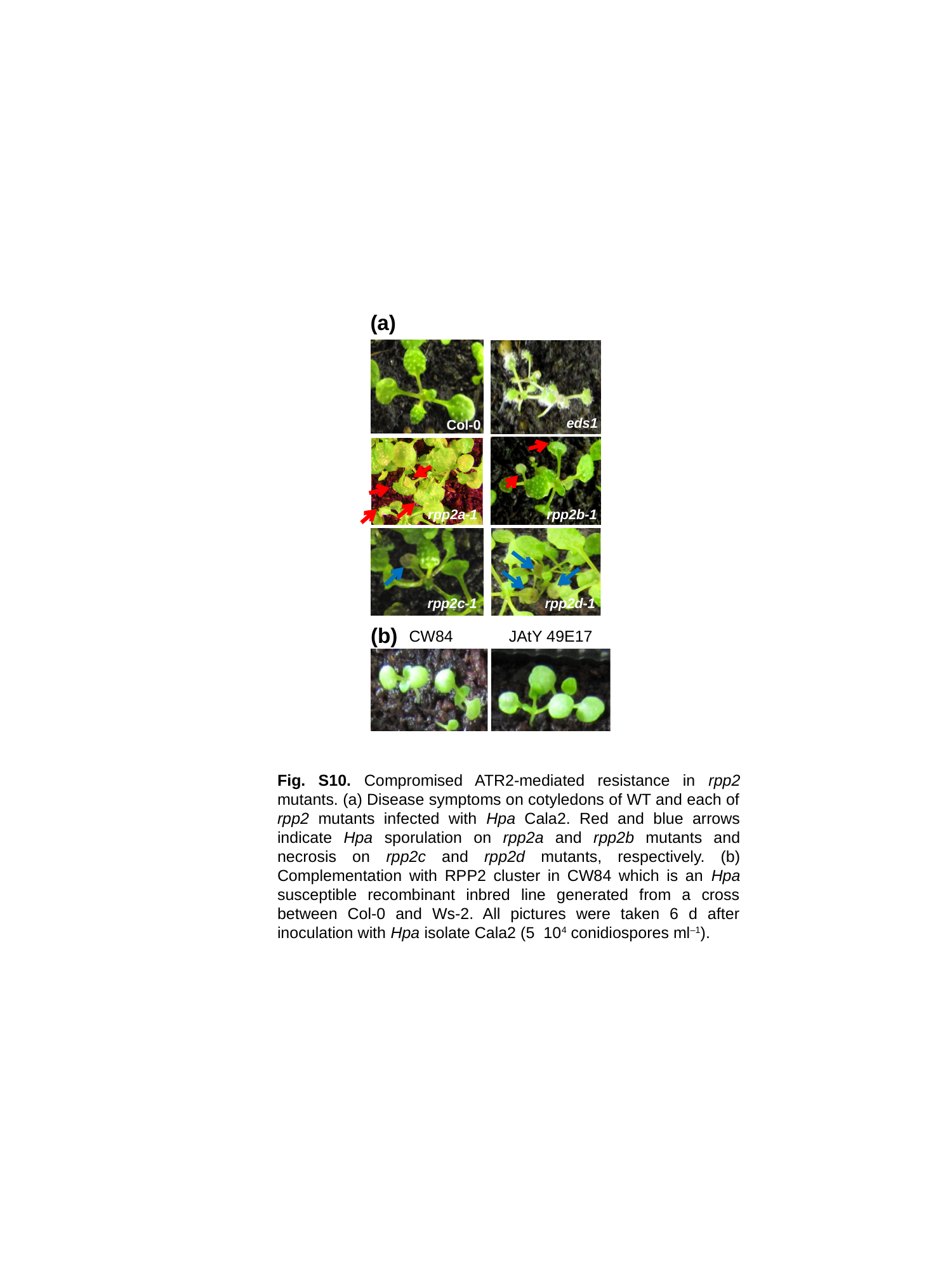

(a)
eds1
Col-0
rpp2b-1
rpp2a-1
rpp2c-1
rpp2d-1
(b)
CW84
JAtY 49E17

### Slide 11
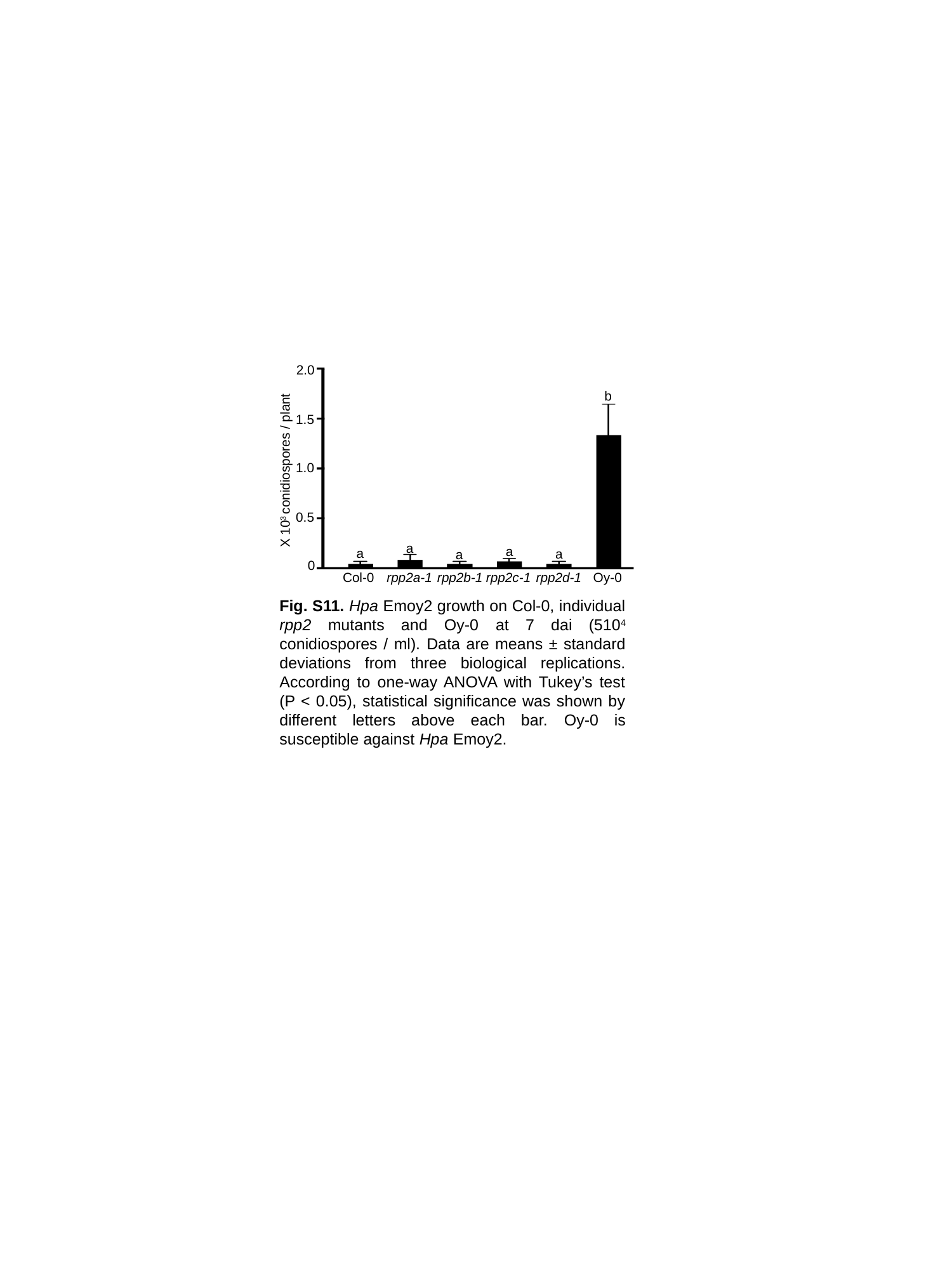

2.0
b
1.5
1.0
X 10^3^ conidiospores / plant
0.5
a
a
a
a
a
0
Col-0
rpp2a-1
rpp2b-1
rpp2c-1
rpp2d-1
Oy-0

### Slide 12
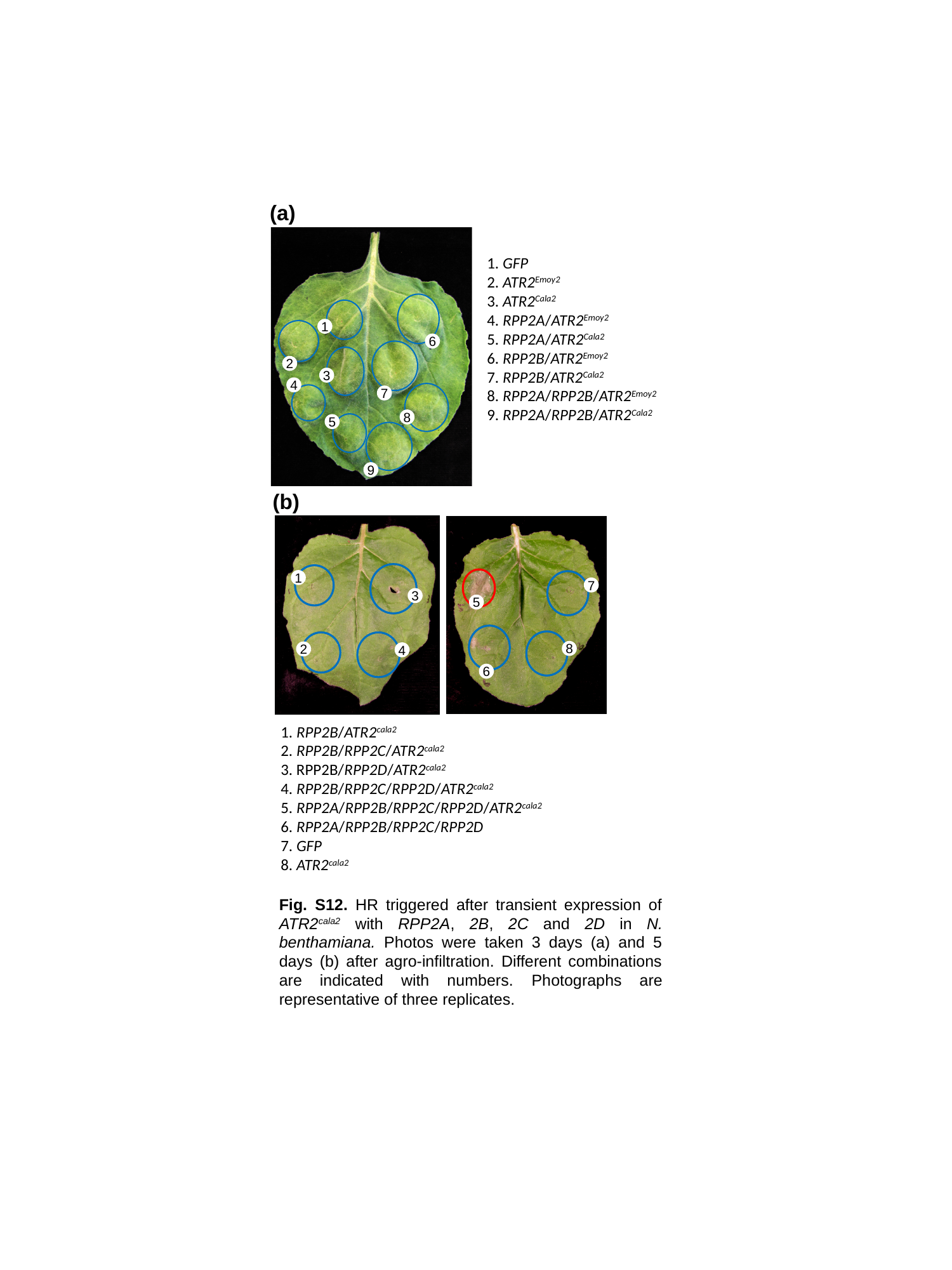

(a)
1. GFP
2. ATR2^Emoy2^
3. ATR2^Cala2^
4. RPP2A/ATR2^Emoy2^
5. RPP2A/ATR2^Cala2^
6. RPP2B/ATR2^Emoy2^
7. RPP2B/ATR2^Cala2^
8. RPP2A/RPP2B/ATR2^Emoy2^
9. RPP2A/RPP2B/ATR2^Cala2^
1
6
2
3
4
7
8
5
9
(b)
1
7
3
5
8
2
4
6
1. RPP2B/ATR2^cala2^
2. RPP2B/RPP2C/ATR2^cala2^
3. RPP2B/RPP2D/ATR2^cala2^
4. RPP2B/RPP2C/RPP2D/ATR2^cala2^
5. RPP2A/RPP2B/RPP2C/RPP2D/ATR2^cala2^
6. RPP2A/RPP2B/RPP2C/RPP2D
7. GFP
8. ATR2^cala2^
Fig. S12. HR triggered after transient expression of ATR2^cala2^ with RPP2A, 2B, 2C and 2D in N. benthamiana. Photos were taken 3 days (a) and 5 days (b) after agro-infiltration. Different combinations are indicated with numbers. Photographs are representative of three replicates.

### Slide 13
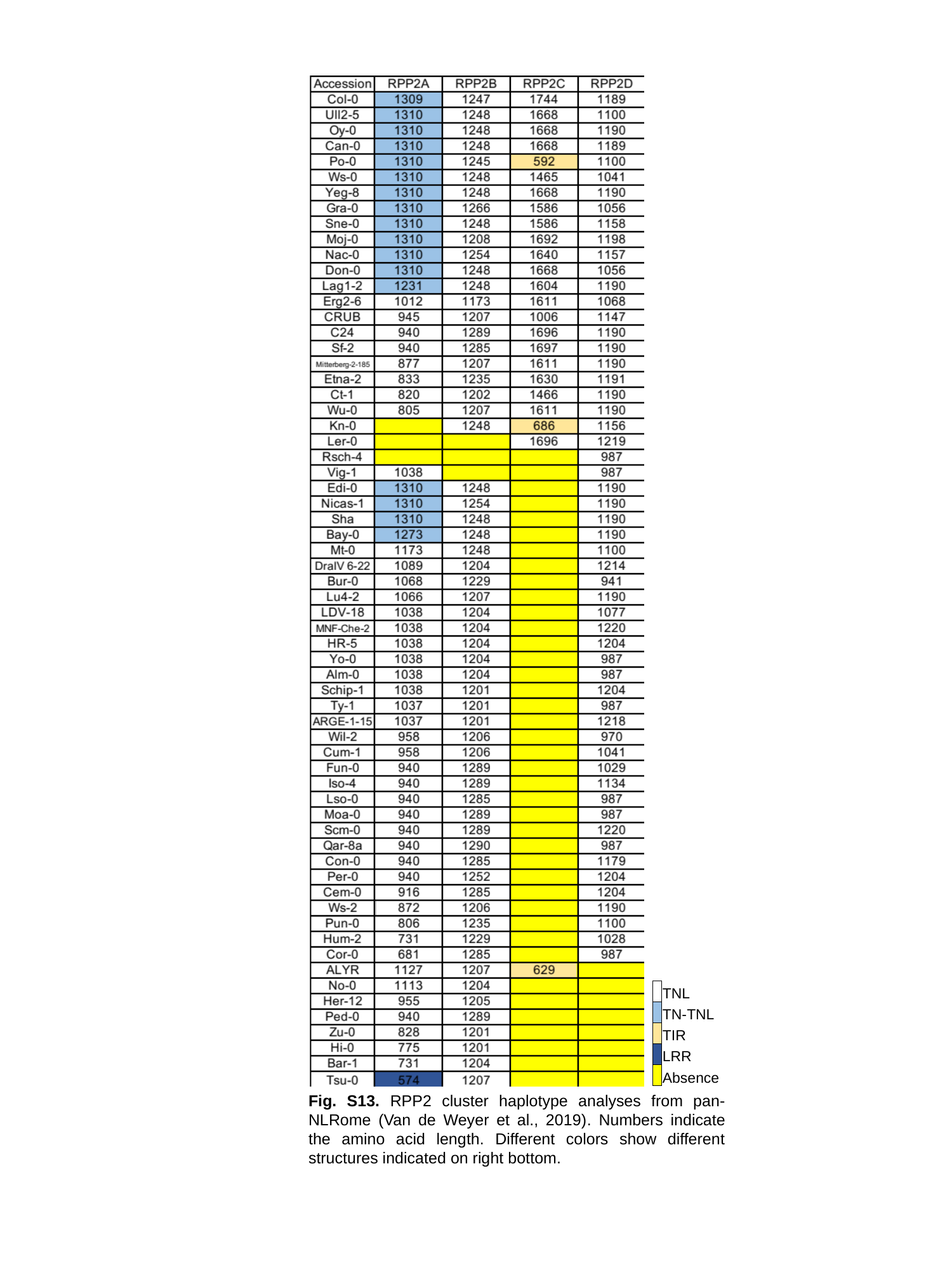

| | TNL |
| --- | --- |
| | TN-TNL |
| | TIR |
| | LRR |
| | Absence |
Fig. S13. RPP2 cluster haplotype analyses from pan-NLRome (Van de Weyer et al., 2019). Numbers indicate the amino acid length. Different colors show different structures indicated on right bottom.

### Slide 14
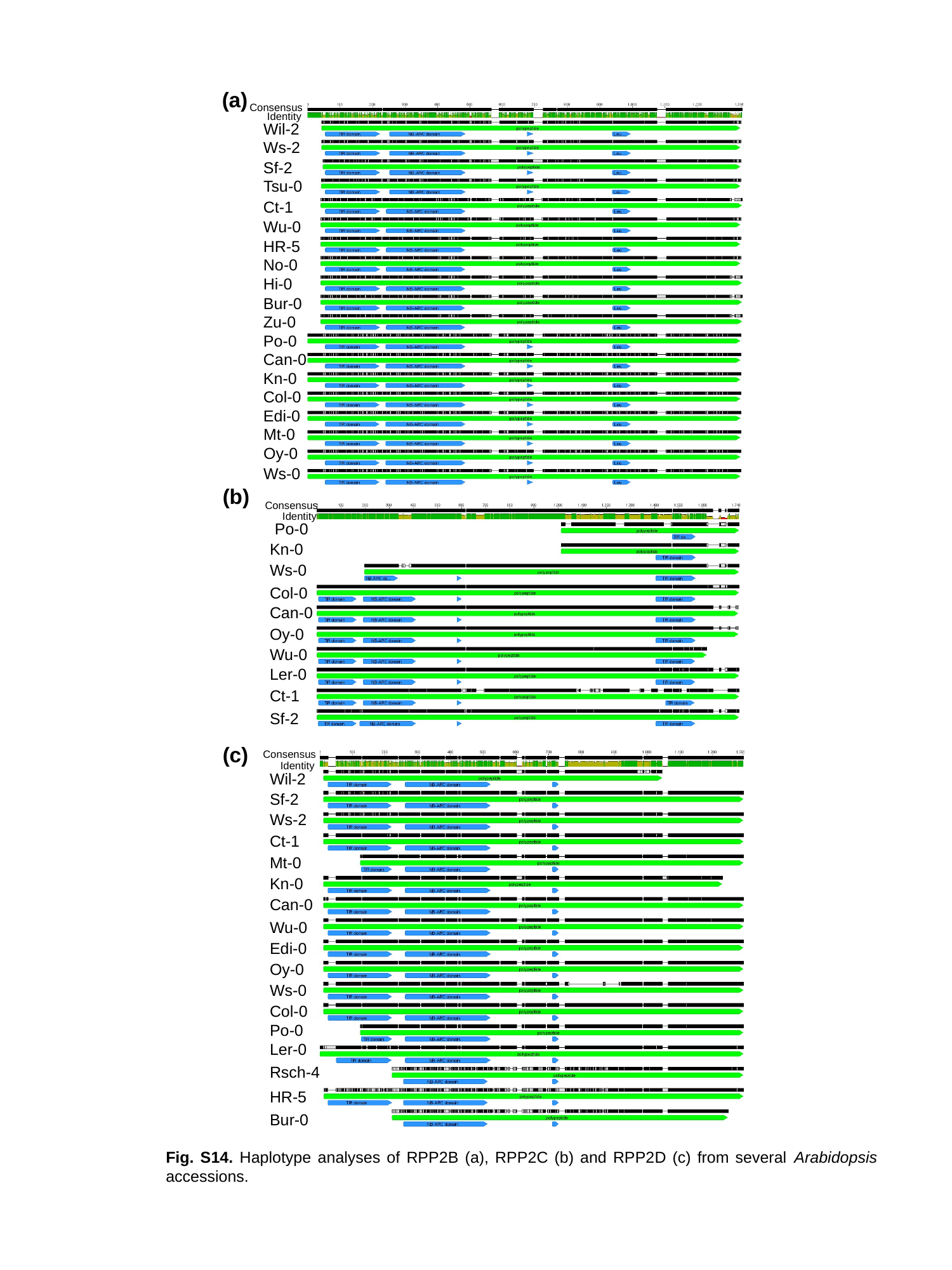

(a)
Consensus
Identity
Wil-2
Ws-2
Sf-2
Tsu-0
Ct-1
Wu-0
HR-5
No-0
Hi-0
Bur-0
Zu-0
Po-0
Can-0
Kn-0
Col-0
Edi-0
Mt-0
Oy-0
Ws-0
(b)
Consensus
Identity
Po-0
Kn-0
Ws-0
Col-0
Can-0
Oy-0
Wu-0
Ler-0
Ct-1
Sf-2
(c)
Consensus
Identity
Wil-2
Sf-2
Ws-2
Ct-1
Mt-0
Kn-0
Can-0
Wu-0
Edi-0
Oy-0
Ws-0
Col-0
Po-0
Ler-0
Rsch-4
HR-5
Bur-0
Fig. S14. Haplotype analyses of RPP2B (a), RPP2C (b) and RPP2D (c) from several Arabidopsis accessions.

### Slide 15
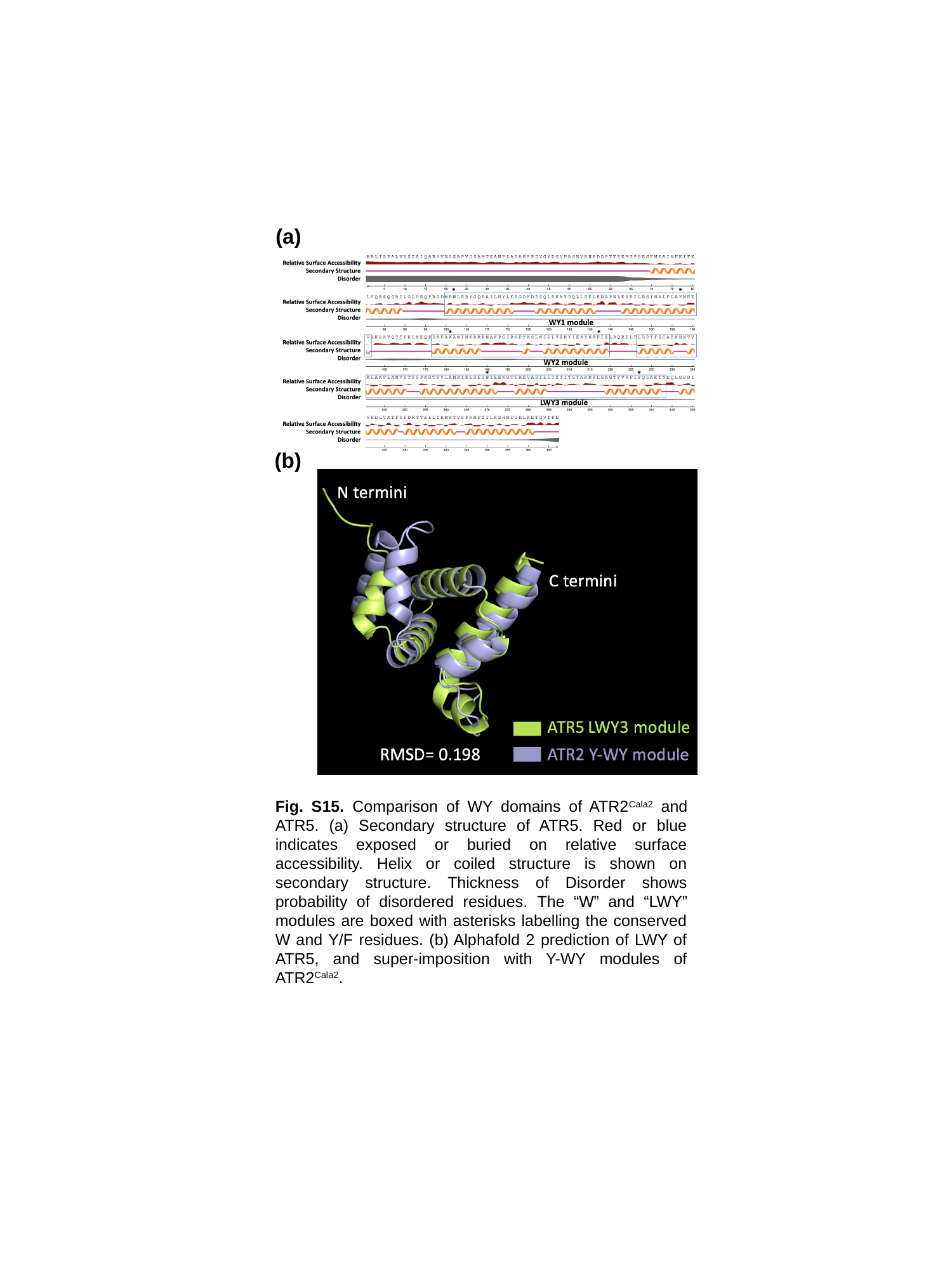

(a)
(b)
Fig. S15. Comparison of WY domains of ATR2^Cala2^ and ATR5. (a) Secondary structure of ATR5. Red or blue indicates exposed or buried on relative surface accessibility. Helix or coiled structure is shown on secondary structure. Thickness of Disorder shows probability of disordered residues. The “W” and “LWY” modules are boxed with asterisks labelling the conserved W and Y/F residues. (b) Alphafold 2 prediction of LWY of ATR5, and super-imposition with Y-WY modules of ATR2^Cala2^.
