## Supplementary material for "ATR2^Cala2^ from *Arabidopsis*-infecting downy mildew requires 4 TIR-NLR immune receptors for full recognition": Methods S1

**Supplemental Materials and Methods**

**Positional cloning of *ATR2^Cala2^***

**Protein gel blot**

Protein was extracted from *Agrobacterium*-infiltrated *N. benthamiana* leaves at 72 dpi as previously described (Kim et al., 2015). Briefly, leaves were harvested, ground in liquid N_2_, and protein was extracted using a buffer (20 mM Tris-HCl, pH7.5, 150 mM NaCl, 1 mM EDTA, 1% Triton X-100, 0.1% SDS, 5 mM 1,4-dithiothreitol (DTT), and protease inhibitor cocktail (Roche). Insoluble debris was pelleted by centrifuging leaf extracts at 15,000 g for 30 min at 4°C. Supernatant from the sample extract was used to elute the samples in SDS-loading buffer. For SDS-PAGE, samples were heated for 10 min at 95 °C for denaturation. After electrophoresis, separated proteins were transferred to PVDF (Merck) membranes for immunoblotting. Membranes were blocked for 2 hours in 5% skimmed milk and probed with horseradish peroxidase (HRP)-conjugated antibodies for overnight at 4°C. Chemiluminescence detection for proteins was carried out by incubating the membrane with developing reagents (SuperSignal West Pico & West Femto).

**Resistance (*R*) gene sequence capture (RenSeq)**

RenSeq experiments were conducted on gDNA freshly extracted from young leaves using the DNeasy Plant Mini Kit (Qiagen, Hilden, Germany) according to the manufacturer’s protocol. For the Illumina MiSeq RenSeq experiments, 2 µg of gDNA were fragmented with the Covaris sonicator (Covaris Inc., MA, USA) using preset 1-kb settings. Fragments were size-selected for fragments longer than 500 bp using Agencourt AMPure XP beads (Beckman Coulter, CA, USA) with 1:0.55 ratio of sheared DNA to beads. Double-stranded cDNA prepared as described above was used directly for library preparation, without shearing or size selection. Illumina MiSeq gDNA and cDNA libraries were prepared using the NEB Next Ultra DNA Library Prep Kit for Illumina (New England BioLabs, Inc., Ipswich, MA, USA) following manufacturer’s instructions. Target capture was carried out using a custom MYcroarray MYbaits kit (MI, USA) and the corresponding protocol. The bait library was designed as described (Van de Weyer et al., 2019). Ten µl of enriched library was PCR-amplified (KAPA HiFi polymerase) up to a quantity of 1 µg in 50 µl reaction volumes using Illumina P5 and P7 primers (Illumina Inc., CA, USA). MiSeq 250-bp paired-end (PE) sequencing was carried out at The Genome Analysis Center (TGAC, Norwich Research Park, UK). Quality control, mapping and SNP calling of Illumina Inc. (San Diego, CA, USA) paired-end sequences were carried out using tools embedded in The Sainsbury Laboratory (TSL) customized Galaxy platform (Maclean and Kamoun., 2012).

**Bioinformatics**

The EnsemblProtist (http://protists.ensembl.org) database were used for *Hpa* genomic contig and EST information*.* Web servers, including InterPro (Quevillon et al., 2005; http://www.ebi.ac.uk/interpro/), and Pfam (Punta et al., 2012; http://pfam-legacy.xfam.org) were used to analyse putative effector proteins, and SignalP V4.0 (Petersen et al., 2011; http://www.cbs.dtu.dk/services/SignalP/) and TMHMM (Krogh et al., 2001; http://www.cbs.dtu.dk/services/TMHMM/) were used to examine each ORF for the presence of a signal peptide and transmembrane helices. Primer designs, and comparison of genomic and full-length cDNA sequences of candidate genes were performed using Geneious Prime.v2015 (Kearse et al., 2012).

**References**

**Bailey K, Cevik V, Holton N, Byrne-Richardson J, Sohn KH, Coates M, Woods-Tör A, Aksoy HM, Hughes L, Baxter L, et al.** 2011. Molecular cloning of ATR5^Emoy2^ from *Hyaloperonospora arabidopsidis*, an avirulence determinant that triggers RPP5-mediated defense in *Arabidopsis. Molecular Plant-Microbe Interactions* **24**(7): 827-838.

**Kearse M, Moir R, Wilson A, Stones-Havas S, Cheung M, Sturrock S, Buxton S, Cooper A, Markowitz S, Duran C, et al.** 2012. Geneious basic: An integrated and extendable desktop software platform for the organization and analysis of sequence data. *Bioinformatics* **28**(12): 1647-1649.

**Kim DS, Kim NH, Hwang BK.** 2015. Glycine-rich RNA-binding protein1 interacts with receptor-like cytoplasmic protein kinase1 and suppresses cell death and defense responses in pepper (*Capsicum annuum*). *New Phytologist* **205**(2): 786-800.

**Krogh A, Larsson B, von Heijne G, Sonnhammer ELL.** 2001. Predicting transmembrane protein topology with a hidden Markov model: application to complete genomes. *Journal of Molecular Biology* **305**(3): 567-580.

**Maclean D, Kamoun S.** 2012. Big data in small places. *Nature Biotechnology* **30**(1): 33-34.

**Petersen TN, Brunak S, von Heijne G, Nielsen H.** 2011. SignalP 4.0: discriminating signal peptides from transmembrane regions. *Nature Methods* **8**(10): 785-786.

**Punta M, Coggill PC, Eberhardt RY, Mistry J, Tate J, Boursnell C, Pang N, Forslund K, Ceric G, Clements J, et al.** 2012. The Pfam protein families database. *Nucleic Acids Research* **40**(D1): D290-D301.

**Quevillon E., Silventoinen V, Pillai S, Harte N, Mulder N, Apweiler R, Lopez R.** 2005. InterProScan: protein domains identifier. *Nucleic Acids Research* **33**(2): W116-W120.

**Van de Weyer AL, Monteiro F, Furzer OJ, Nishimura MT, Cevik V, Witek K, Jones JDG, Dangl JL, Weigel D, Bemm F.** 2019. A species-wide inventory of NLR genes and alleles in *Arabidopsis thaliana*. *Cell* **178**(5): 1260-1272.

**Woods-Tör A, Studholme DJ, Cevik V, Telli O, Holub EB, Tör M.** 2018. A suppressor/avirulence gene combination in *Hyaloperonospora arabidopsidis* determines race specificity in *Arabidopsis thaliana*. *Frontiers in Plant Science* **9**: 265.
